# Dynamic resource allocation orchestrates physical simulation in the human brain

**DOI:** 10.64898/2026.06.21.732202

**Authors:** Ling Long, Qiyue Wang, Qianli Yang, Le Chang

## Abstract

Humans can flexibly predict physical events by internally simulating object dynamics in the brain—a capacity lacking in current AI systems. Using a ball collision paradigm with visual occlusion combined with multimodal neuroimaging (fMRI/MEG), we uncover a spatiotemporally organized neural architecture for physical simulation. fMRI reveals hierarchical spatial segregation: higher-order cortical regions encode relational physical variables, distinct from object-specific features encoded in the lower-order sensorimotor regions. MEG uncovers two temporally distinct neural processes: a real-time simulation tracking the evolving state of the object, in alignment with collision dynamics, and an early predictive signal anticipating collision occurrence (∼700 ms before contact). We propose that this early predictive control mechanism dynamically allocates cognitive resources during simulation. This is formalized by a Dynamic Resource Intuitive Physics Engine (IPE) model, which captures both behavioral data and the dual neural timescales by optimizing accuracy-cost tradeoffs. Crucially, this framework predicts early encoding of another adaptive control variable separate from collision occurrence, as evidenced by both MEG and pupillary responses. These findings reveal how the brain achieves efficient physical inference through hierarchically organized predictive control that dynamically allocates cognitive resources.

## Introduction

Humans possess a remarkable ability to understand and predict the dynamics of physical scenes, enabling them to act preemptively in complex, real-world situations. During a soccer match (Fig. 1A), for example, a player continuously predicts the ball’s future trajectory based on its current motion state and simulates how different kicking actions would alter its path, thereby enabling precise motor execution. Such predictive simulation allows for fluid, adaptive behavior in dynamic environments. In contrast, current large vision-language models—despite their remarkable progress in recognition and semantic reasoning—exhibit limited physical world understanding. They often struggle with physical object properties, physical scene understanding, and dynamic interactions, revealing their limitations in intuitive physical reasoning^1^.

**Figure 1.**
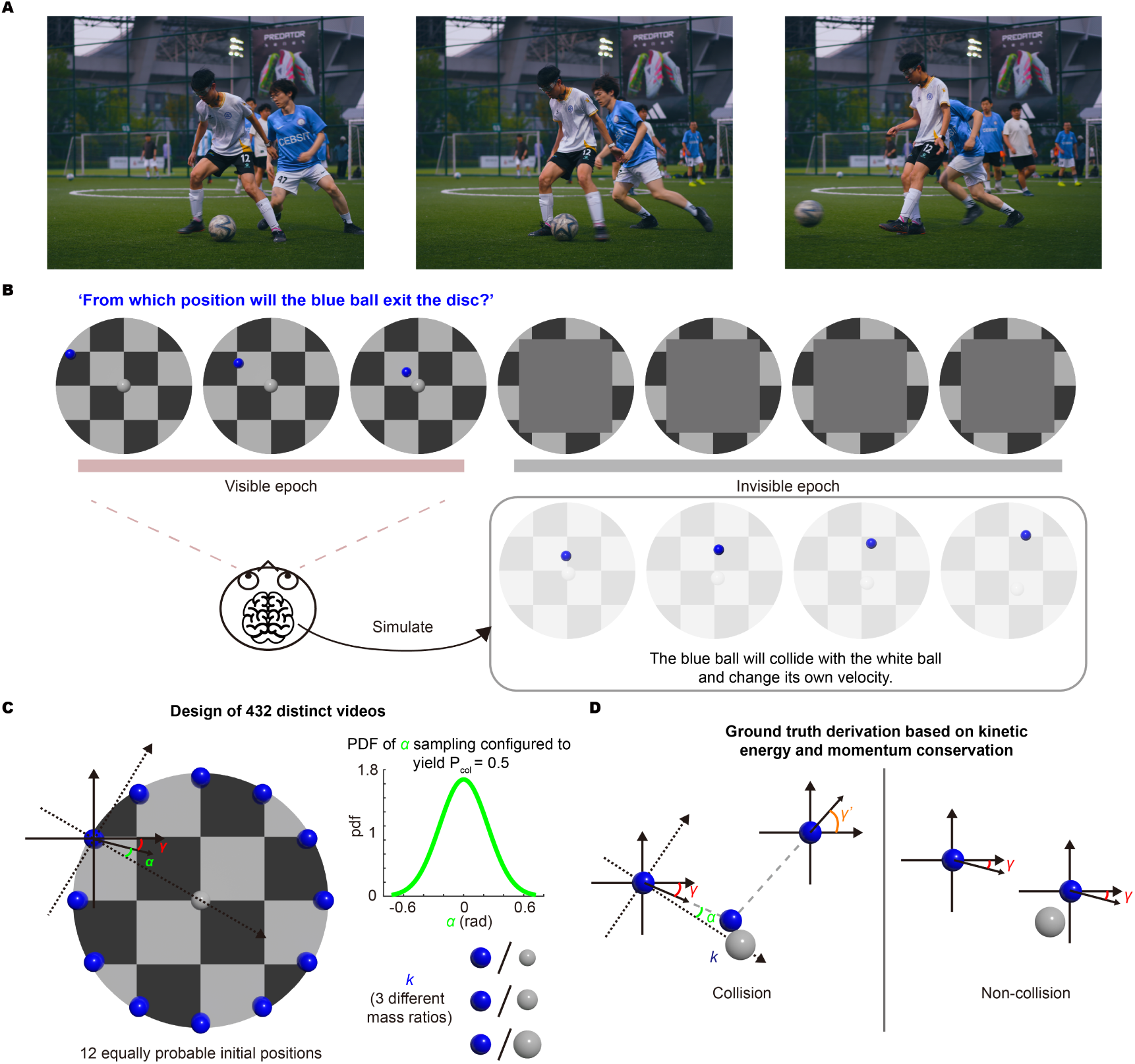
How the brain simulates physics: a parametric ball collision scenario. (A) Humans possess a deep understanding of physical scene dynamics. In this soccer scene, the player in white continuously predicts the ball’s future trajectory based on its current motion state and simulates how different kicking actions would alter its path. At the same time, the player in white mentally simulates how the blue player’s kick would alter the ball’s trajectory, based on the motion of the ball and the body posture of his opponent, thereby planning his defensive strategy. These photographs were taken by the authors with written informed consent from the individuals shown. (B) A ball-collision physical scenario was designed to investigate the neural mechanisms underlying internal simulation of physical scenes. Participants viewed videos showing collisions between two balls, which were occluded shortly before the collision. Their task was to predict exit position of the blue ball by internally simulating the collision process. (C) A total of 432 videos were created. The blue ball appeared with equal probability at one of 12 discrete positions, evenly spaced around the disc. The interaction between the blue and white balls was governed by the relative angle α between the blue ball’s initial motion direction and the line connecting the centers of the two balls. This angle was sampled from a Gaussian distribution calibrated to yield collisions in approximately 50% of trials. Additionally, three mass ratios between the two balls were included (k = m_blue / m_white: 3.375, 1, 0.422). (D) Based on the relative angle α, trials were naturally divided into collision and non-collision trials. The post-collision motion directions were computed using classical Newtonian mechanics, providing a ground truth for the participants’ predictions.

Physicists address this challenge by formulating analytical equations that precisely describe the evolution of physical systems. In contrast, the biological brain acquires this predictive capacity through experience. From early infancy, humans gradually internalize the statistical regularities of physical interactions, ranging from recognizing that an apple resting on a table remains supported to anticipating the trajectory of a kicked soccer ball^2–6^. While simpler rules may emerge early during development, mastering more complex dynamics—such as collisions or unstable configurations—requires prolonged interaction with the physical world. Nevertheless, once acquired, this internalized knowledge enables humans to predict physical scene dynamics with remarkable speed and robustness, while generalizing flexibly across contexts.

This intuitive predictive capability is thought to rely on an internal mental model within the human brain^7–18^. This model is capable of constructing dynamic mental representations of physical systems and forecasting future outcomes. Neuroscientific evidence from fMRI implicates a fronto-parietal network^15^ in supporting such simulations, where multivariate activation patterns across this network collectively encode abstract properties such as mass and stability^16–18^. While existing studies have established a coarse-grained correspondence between the fronto-parietal network and physical simulation, this framework fails to account for both the structural and functional complexity of the human brain, as well as the rich spatiotemporal dynamics of real-world physical scenes where multiple objects interact over time. Consequently, the fine-grained neural mechanisms underlying physical simulations remain poorly understood.

This limitation presents a fundamental challenge for developing neural network models that aim to emulate human physical reasoning^1,8,19–24^, as even state-of-the-art computer vision models trained on natural videos fail to match the brain’s predictive abilities in physical scenarios^1,19,20^. However, Intuitive Physics Engines (IPEs)—a class of iterative probabilistic generative models—have shown considerable explanatory power for human physical reasoning behavior^8,24^. Crucially, a key feature of such probabilistic simulation is the presence of uncertainty, stemming not only from sensory ambiguity but also from internal noise introduced at each step of simulation. In real-world environments, where multiple objects interact via complex nonlinear dynamics, even small amounts of noise can amplify into large deviations in simulated outcomes, making effective management of uncertainty essential for biological physical reasoning. Yet it remains unclear whether this uncertainty operates as a fixed computational constraint or a dynamic parameter modulated by higher cognitive functions (e.g., attention). Clarifying this distinction would advance our understanding of uncertainty management in neural physical reasoning, creating opportunities to align computational models with biological principles.

To address these questions, we developed a naturalistic yet parameterized ball collision paradigm that captures dynamic physical interactions while allowing systematic manipulation of key variables. Through temporally controlled visual occlusion, this paradigm effectively disentangled internal physical simulation from online perceptual processing. Using a combination of behavioral, functional magnetic resonance imaging (fMRI), and magnetoencephalography (MEG) measurements, we investigated both the spatial organization and temporal dynamics of physical simulation in the human brain.

Our findings reveal a highly structured neural architecture for physical simulation that diverges from the traditional views. Spatially, distinct physical variables are represented in anatomically segregated cortical regions organized along the sensorimotor-to-executive hierarchy gradient. Temporally, we observed dual neural timescales: (1) rapid predictive responses to critical physical events (e.g., collision) and (2) real-time simulations tracking the evolving state of object, in alignment with collision dynamics. These observations necessitate an extended IPE framework in which the brain dynamically regulates resource allocation to optimize the accuracy-cost tradeoff during multi-timescale simulation.

## Results

We designed a visual paradigm with parameterized control over key variables, simulating elastic collisions between two balls, and presented it to human participants as short video clips. Each video began with a stationary white ball positioned at the center of a disc, while a blue ball started from a random position on the boundary of the disc and moved at a constant speed in a randomized direction. At a predetermined time point, prior to the anticipated collision, the two balls were occluded. The participants were then asked to predict where the blue ball would eventually exit the disc (Fig. 1B).

A total of 432 video stimuli were designed. At the beginning of each video, the blue ball was positioned in one of 12 equidistant locations spaced uniformly around the disc’s boundary (Fig. 1C). The initial motion direction of the blue ball was quantified using two coordinate systems. In the world coordinate system, it was represented by the absolute angle *γ*. In the relative coordinate system—defined by the axis connecting the centers of the two balls (the inter-ball axis) and its normal vector—it was represented by the relative angle *α*. The mass ratio between the two balls was varied (denoted by *k,* three levels in total), with their masses being proportional to their volumes. Prior to the main experiment, all participants watched a 1-minute preview of collision videos in which the full trajectories were visible, randomly selected from the complete stimulus set, to become familiar with the task structure and the underlying physical principles.

Throughout this paper, the term “ground truth” refers to the blue ball’s trajectory for a given trial, computed analytically from Newtonian mechanics (Fig. 1D). If a collision occurred, the motion direction of the blue ball was updated accordingly (=final motion direction, denoted by *γ′*); otherwise, its original motion direction *γ* was maintained. The collision detection threshold was crucial—when the blue ball’s initial motion direction passed a critical value, the scenario transitioned sharply from non-collision to collision, resulting in a dramatic change to the required calculations. While physical variables such as the relative angle *α* and the mass ratio *k* were present in all trials, they only became relevant in collision scenarios. This binary distinction critically affects the participants’ simulation strategies. Elucidating the neural mechanisms underlying collision detection was therefore a central aim of the study.

Based on this experimental design, we conducted a behavioral study (n = 16 participants), an fMRI study (n = 11 participants), and two MEG studies (n = 17 participants for Experiment 1 and n = 12 participants for Experiment 2). All studies shared the same core task where participants predicted the post-collision trajectory of the blue ball, though response granularity varied between precise angle reporting and coarse categorizations.

### Human predictions of collisions adhere to Newtonian physics

Predicting the motion of the blue ball in non-collision trials seems to be a straightforward task. The intuitive physical principle that objects maintain their state of motion in the absence of external forces has been shown to develop in human infants as young as six months old.^2^ However, can human participants accurately predict the motion of the blue ball following a collision? This question is particularly intriguing given that collision dynamics often involve nonlinear computations, where changes in velocity and direction arise from complex interactions between mass, momentum, and contact geometry.

In the behavioral study, participants were instructed to click on the position along the edge of the disc where they predicted the blue ball would exit, once the disc’s color changed from red to green (Fig. 2A). The clicked positions were converted into angular values within a world coordinate system centered at the disc’s origin (Fig. 2B). We computed the mean absolute error (MAE, in radians) between each participant’s predicted exit angle and the ground truth angle, analyzed separately for collision and non-collision trials. The results showed that in the non-collision trials, participants’ predictions had very low MAE, indicating close alignment with the ground truth. In contrast, collision trials yielded higher MAE and greater between-participant variability (Fig. 2C, open circles). Nevertheless, all participants performed significantly above chance in both collision and non-collision trials (p < 0.001 in both, 20,000 random shuffles).

**Figure 2.**
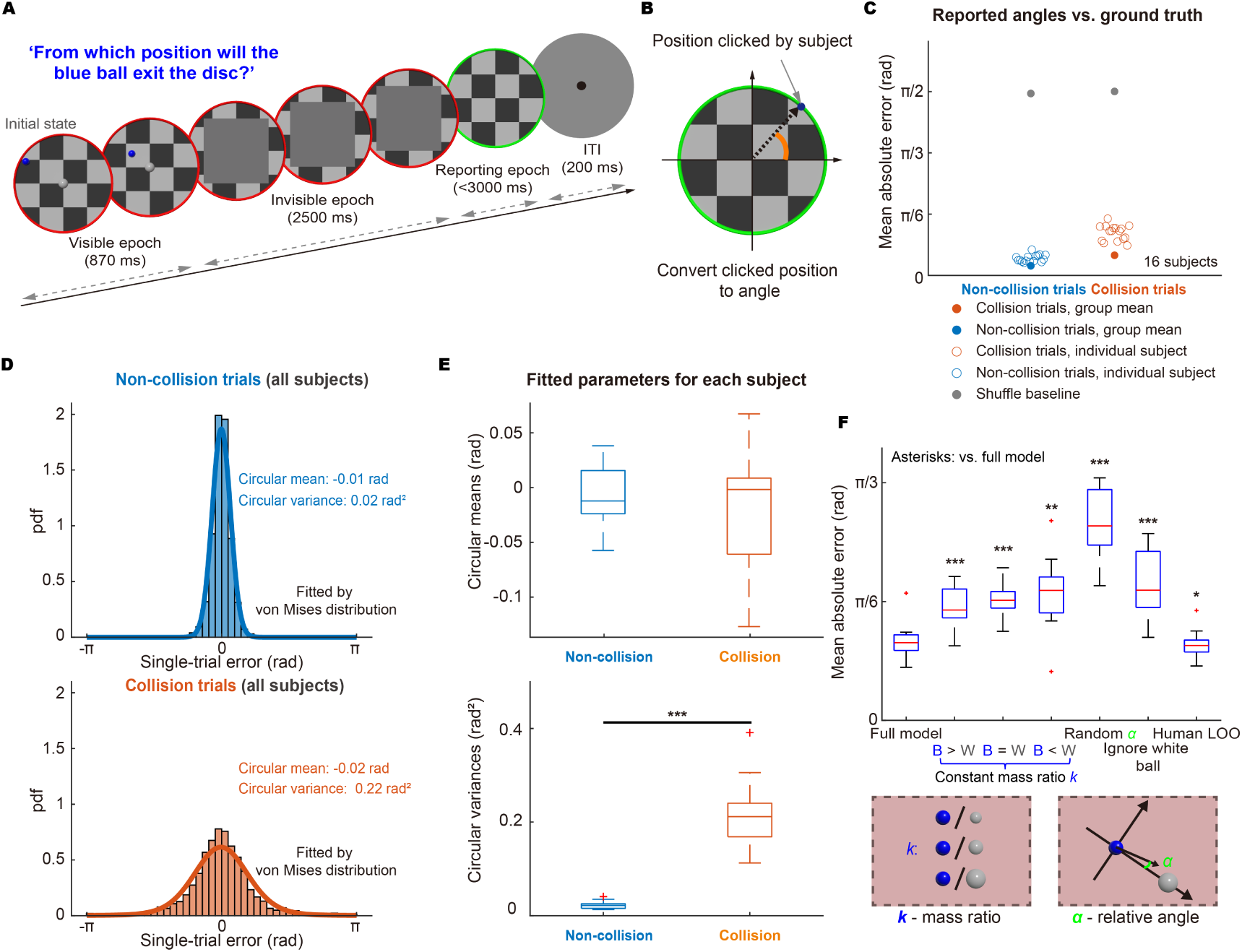
Behavioral results. (A) Participants viewed videos of two-ball collisions (432 trials; collision probability = 0.5). The two balls were occluded prior to anticipated collision. Participants were asked to predict the post-collision exit point of the blue ball from the disc, indicated by a color change of its edge from green to red. (B) The position of the participant’s click was converted into an angle for later analyses. (C) Mean absolute error (MAE; mean of the minimal absolute differences in exit angle, in radians) between participants’ predictions and the ground truth (derived from Newtonian mechanics; 16 participants). Non-collision trials demonstrate consistently low MAE for both individual participants (blue, open circles) and the group average (blue, solid circle, averaged across participants for each trial). In collision trials, individual participants (orange, open circles) exhibit higher MAE, whereas the group average (orange, solid circle) maintains a low MAE (0.17 rad). Gray dots indicate the shuffle baseline. (D) The von Mises distribution was fitted to the prediction errors aggregated across all participants (R^2^ = 0.95 and 0.96 for collision and non-collision trials, respectively). (E) For each participant, prediction errors were fitted with the von Mises distribution separately for the collision trials and for the non-collision trials. No significant difference in the circular mean was observed between the two trial types, but the circular variance differed significantly (*** = p < 0.001, two-tailed Student’s t-test). (F) Three types of perturbation were made to the parameters of the full Newtonian mechanics model: 1) fixing the mass ratio k across conditions; 2) randomizing the relative angle α; 3) ignoring the white ball. Compared with the full model, these perturbations yielded significantly higher MAE between model outputs and participants’ predictions. A human noise floor was estimated using a leave-one-subject-out procedure: for each participant, the mean absolute error (MAE) of their responses was computed relative to the circular mean of all other participants for the same stimulus. Asterisks indicate significant differences relative to the full model, assessed using two-sided Wilcoxon signed-rank tests (* = p < 0.05, ** = p < 0.01, *** = p < 0.001).

However, when averaging predictions across participants for each of the 432 scenes, the group-averaged predictions showed low MAE relative to the ground truth in both conditions (MAE = 0.17 rad for collision trials and 0.08 rad for non-collision trials) (Fig. 2C, solid circles). To explain this effect, we further examined the distribution of single-trial prediction errors across all participants, analyzed separately for collision and non-collision trials. Both distributions were well fit by a von Mises distribution (collision trials: *R²* = 0.95; non-collision trials: *R²* = 0.96) (Fig. 2D). While the circular mean of the prediction errors was near zero in both conditions, the circular variance was considerably larger in the collision trials than in the non-collision trials. Fitting the prediction error distribution for individual participants validated these results (Fig. 2E; circular mean: t = 0.6108, p = 0.5459; circular variance: t = 11.0945, p < 0.001; two-tailed Student’s t-tests, collision vs. non-collision). Thus, this pattern can be explained by greater inter-individual variability in collision trials and the absence of a systematic directional bias, such that individual errors tended to cancel out when averaged across participants.

Despite substantial individual variability in collision trials, we observed systematic dependencies of prediction errors on key physical variables. Specifically, the MAE exhibited a parametric relationship with both relative angle *α* and mass ratio *k* (Fig. S1), displaying Gaussian-like patterns centered at *α* = 0 with distinct profiles across mass ratio conditions. The structured covariation between errors and physical variables suggests that participants’ judgments were systematically influenced by task-relevant physical information.

To directly test this possibility, we generated predictions derived from both the full Newtonian mechanics model and perturbed models featuring systematically distorted physical variables (Fig. 2F). The human noise floor, computed using a leave-one-subject-out procedure, is also shown for comparison. The MAE between the full model and participants’ predictions was significantly lower than that of all perturbed models (Wilcoxon signed-rank tests, two-sided; four perturbations: p < 0.001, one perturbation p < 0.01), but was slightly higher than the human noise floor (p < 0.05).

### Spatially segregated representations of physical variables across cortical hierarchies

We next used fMRI to identify brain regions representing task-relevant physical variables during the ball-collision task, such as relative angle α and mass ratio k. In the full model, these variables jointly determine the directional change of the blue ball from *γ* (initial velocity direction) to *γ′* (final velocity direction). The fMRI experiment used an event-related design in which the 432 unique collision scenes were presented as randomized individual trials. To dissociate neural activity related to participants’ simulation outcomes (specifically, the predicted final exit point of the blue ball) from motor-related signals, we adapted the reporting procedure from the behavioral experiment. In this task, the disc border was partitioned into two semicircular halves (red and green), and participants were instructed to indicate which half of the border the blue ball would reach (Fig. 3A). Responses were initiated upon appearance of a reporting cue (red and green arrows), with each color systematically mapped to the participant’s right index or middle finger.

**Figure 3.**
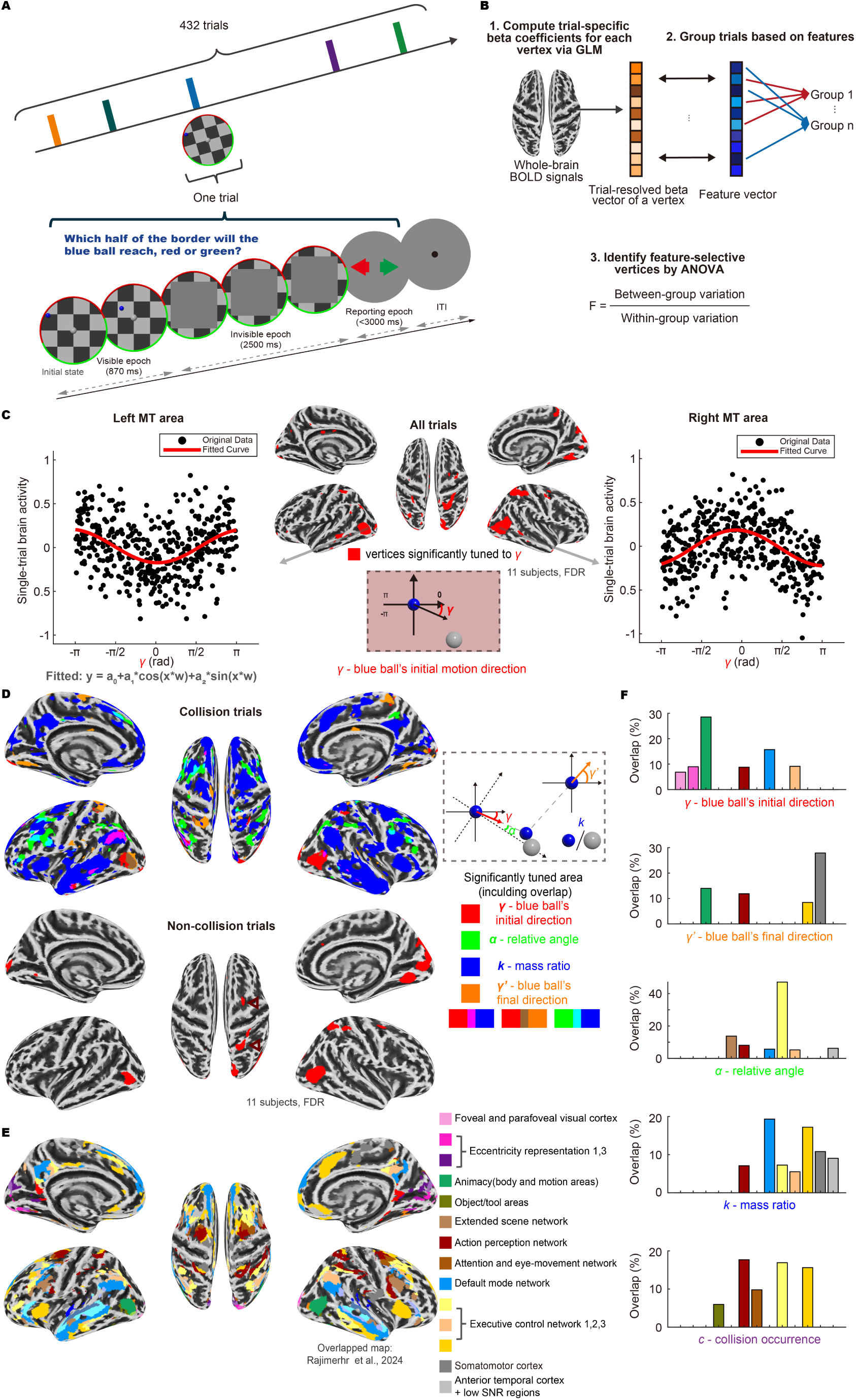
Functional segregation and hierarchical organization within the physical simulation network. (A) Participants in the fMRI experiment indicated whether the blue ball would reach the red or green semicircular arcs, which were randomly rotated in each trial while equally dividing the disc into two halves. (B) Analysis procedure. Trial-specific beta coefficients are computed using a GLM with separate regressors per trial. Trials are then grouped according to physical variables of the dynamic scene, and a one-way ANOVA is performed to identify vertices with significant tuning. (C) The whole-brain search reveals that bilateral MT regions exhibit strong tuning to the initial motion direction γ of the blue ball (n = 11 participants, p < 0.05, FDR correction, same below). Beta vectors from all vertices in the same MT region are averaged, and their relationship with γ is plotted. The tuning curves of the left and right MT regions exhibit opposite trends, consistent with their preference for processing contralateral visual fields. For example, the right MT region responds more strongly to motion in the left visual field, as a blue ball starting in the left visual field tends to move rightward (i.e., γ ≈ 0). (D) All trials in the physical simulation condition are categorized according to whether a collision occurs. For each physical variable, vertices with significant tuning (p < 0.05) are projected onto the inflated cortical surface. Color coding: red and orange for the initial γ and final γ′ motion direction of the blue ball, blue for the mass ratio k, and green for the relative angle α. Significant tuning to two variables is represented by a mixed color as illustrated. No vertices were found to be tuned to three variables simultaneously. Within the bottom panel, the upper red triangle indicates the right superior frontal cortex, while the lower triangle denotes the right superior parietal cortex. (E) The significantly tuned regions in collision trials overlaid on a functional cortical parcellation derived from naturalistic movie-watching fMRI data (Rajimehr et al., 2024; 24 networks). Color codes correspond to the functional networks (right). (F) Spatial overlap (Jaccard Index, JI) between physical variable-tuned regions and movie-derived functional networks (only clusters with JI > 0.05 shown). Variables: blue ball’s initial motion direction γ, blue ball’s final motion direction γ′, relative angle α, mass ratio k and collision occurrence c.

In the fMRI experiment, each of the 432 trials presented a unique scene with parametrically varied physical variables (*α, γ,* etc.). To identify brain regions sensitive to these variables, we conducted a whole-brain surface-based univariate ANOVA on vertex-wise responses (Fig. 3B). The underlying assumption is that if a vertex encodes a given variable, its response dissimilarity should increase with parametric distance between values. To test this, we discretized the variable values into bins and, for each vertex, used F-statistics to quantify whether its responses varied systematically across them. The analysis pipeline began with single-trial response modeling using a general linear model (GLM) approach, where each trial served as an independent regressor^25,26^. This yielded vertex-specific beta coefficients reflecting trial-wise activation. For each physical variable of interest, a feature vector was created to represent parametric variations across trials. Beta coefficients were then binned based on corresponding feature values, enabling the computation of F-statistics to quantify vertex-wise tuning strength for each variable. Finally, vertex-wise F-statistics were assembled into whole-brain maps.

Using this method, we first conducted a whole-brain search to identify regions selectively tuned to the blue ball’s initial motion direction, *γ*. The bilateral MT areas emerged as showing the highest F-statistics (all trials from 11 participants, FDR corrected, *p* < 0.05). Subsequent tuning curve analysis revealed a clear contralateral visual field bias in both MT regions (Fig. 3C): the left MT exhibited preferential tuning to motions in the right visual field, and vice versa. This finding is consistent with both human fMRI studies and macaque electrophysiological recordings targeting the homologous MT region^27,28^, validating our whole-brain univariate ANOVA method for detecting neural tuning to physical variables.

For each physical variable, whole-brain tuning maps were generated, with significantly tuned regions color-coded by variable. Results from collision and non-collision trials were visualized separately to reveal differential spatial distributions (Fig. 3D).

While the blue ball’s initial motion direction *γ* was consistently represented in the MT regions across both conditions, non-collision trials additionally engaged right superior parietal cortex (spatial working memory^29,30^) and right superior frontal cortex (maintenance and manipulation of spatial representations^31^), suggesting sustained mental tracking of the blue ball’s trajectory during occlusion (triangles in Fig. 3D, bottom). To further address the possibility that participants adopted markedly different viewing strategies across trial types, we performed whole-brain multivariate decoding of initial motion direction (*γ*). Reliable decoding was observed within both collision and non-collision trials, with accuracies significantly above chance. Moreover, classifiers trained on one trial type generalized successfully to the other (collision → non-collision and non-collision → collision), with no significant difference between cross-trial-type and within-trial-type decoding performance (Fig. S2A). These results indicate that neural representations of initial motion direction are broadly consistent across conditions, helping to mitigate concerns about potential confounds arising from differences in viewing strategies.

Crucially, significant representations of the relational physical variables underlying collision inference (i.e., *α* and *k*) were only observed in the collision trials (Fig. 3D, top). However, this apparent absence in non-collision trials should not be interpreted as a complete lack of neural information. Indeed, complementary multivariate decoding analyses revealed reliable information about relative angle (*α*) in non-collision trials at the whole-brain level, as well as within regions identified as *α*-selective in collision trials (Fig. S2B). These results suggest that relative-angle information is represented in both conditions, but is less readily detectable in non-collision trials in the univariate analysis.

Furthermore, the representations of different variables showed minimal spatial overlap (overlapping representations are indicated by mixed colors in Fig. 3D), with the number of vertices selective for two or more variables being significantly lower than expected by chance (permutation test, p = 0.020; Fig. S3). This finding suggests that frontoparietal and temporal areas collectively form a physical simulation network that is anatomically distributed yet functionally segregated, where distinct sub-networks preferentially encode specific physical variables.

Our findings align with Fischer et al.’s localization of a frontoparietal physical simulation network^15^, despite methodological differences: whereas their task-based contrasts identified regions activated during block-stacking collapse prediction, our encoding revealed parametric tuning to physical variables in elastic ball collisions. Notably, while the temporal lobe showed no significant activation in traditional contrast analyses comparing collision and non-collision trials (Fig. S4A), explicit encoding modeling revealed its selective representation of mass ratios (Fig. 3D, top), suggesting that conventional activation analysis may overlook parametric physical representations in these regions.

On the representation map derived from the collision trials, the initial and final motion directions of the blue ball (*γ* and *γ’*) were primarily localized in the occipital-temporal-parietal regions, whereas relational physical variables (*α* and *k*) occupied more anterior frontal-parietal regions. To quantify this hierarchical dissociation, we aligned the representation map of each physical variable with Rajimehr et al.’s naturalistic movie-based parcellation (Fig. 3E)^32^. The resultant overlap analysis (Fig. 3F,only functional clusters with a Jaccard Index > 0.05 are shown; see Fig. S4D for full results) revealed that object-specific physical variables (*γ* and *γ′*) predominantly engaged sensory and motor areas (dark green and gray), whereas relational variables (*α* and *k*) preferentially recruited high-level cognitive regions including executive control and default mode networks (yellow and dark blue).

Similar to relational variables, collision-specific activations (contrasting collision and non-collision trials) strongly engaged the executive control networks 1 & 3 (Fig. 3F, Fig. S4B, C). The overlap between regions encoding these abstract relational variables and executive control networks suggests a functional linkage to cognitive control mechanisms, which may modulate physical inference to ensure flexibility.

### Neural dynamics of physical simulation

In spite of a systematic characterization of the spatial patterns of neural representations, their temporal dynamics are elusive due to the low temporal resolution of fMRI data. We addressed this limitation by employing magnetoencephalography (MEG), which provides millisecond-level temporal resolution to better capture these processes. The MEG experiment replicated the fMRI paradigm while adding a background pattern control condition (Fig. 4A). In this control, the white ball served as a non-interactive texture, creating scenarios where the blue ball would continue its motion without collision. Although the video stimuli were identical prior to occlusion, the two conditions required distinct internal simulations: predicting physical interactions in the experimental condition and predicting uniform motion in the control condition. We maintained a trial-by-trial structure with block-wise context cues (physical simulation or background pattern) to ensure task clarity. Eye movements were also monitored during the MEG experiment, allowing us to assess whether collision and non-collision trials were associated with systematically different gaze-tracking behavior within the physical simulation condition (Fig. S5). No reliable differences in gaze tracking patterns were observed between these trial types.

**Figure 4.**
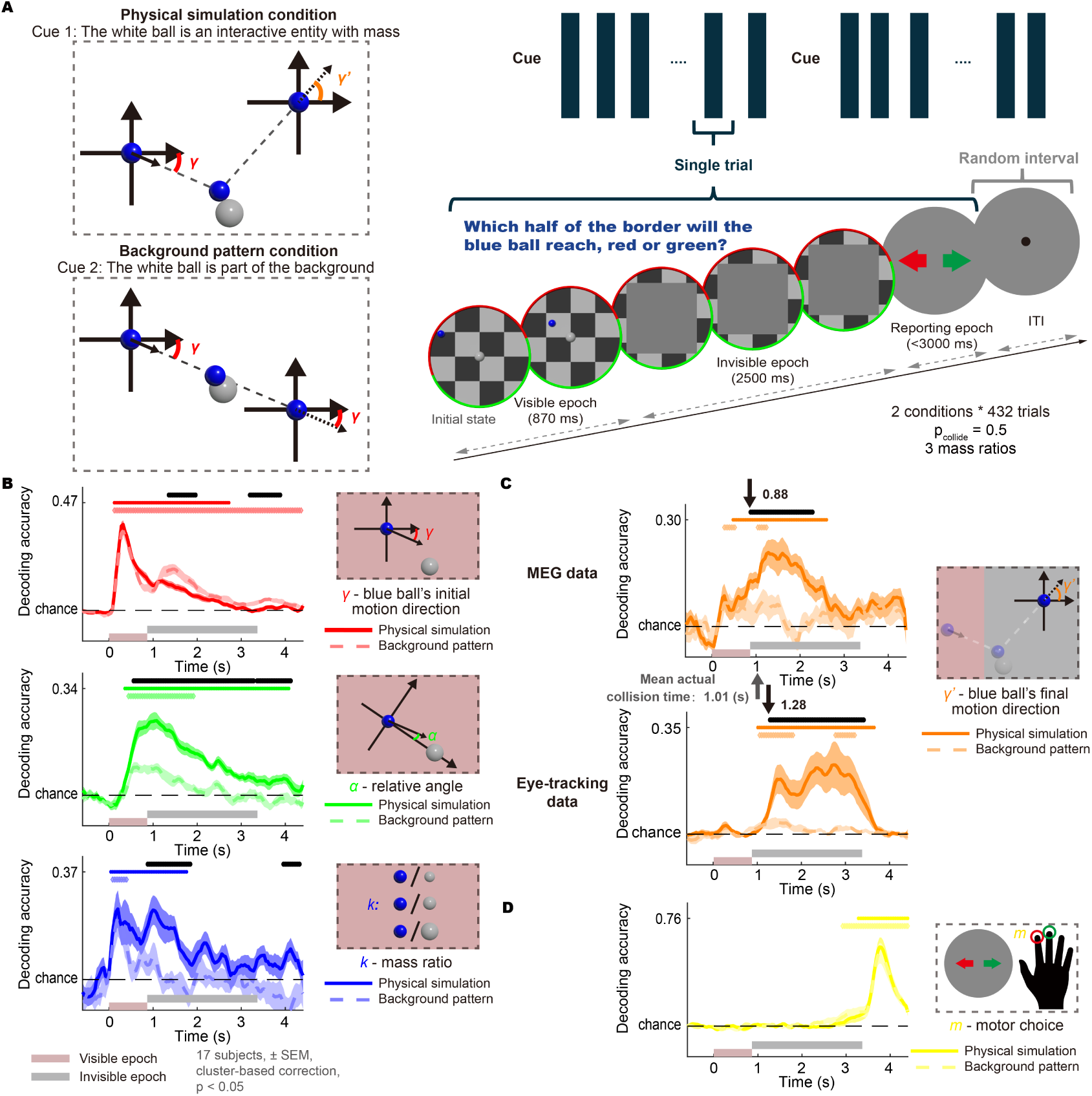
Neural dynamics of different physical variables. (A) The MEG experiment included two distinct stimulus conditions: a physical simulation condition and a background pattern condition (as a control). In the control condition, the white ball was presented as a static background pattern without mass, whereas in the physical simulation condition, it was presented as an interactive object with mass. The visible parts of the scenes were identical in both conditions; however, different processes need to be simulated during the invisible epoch. The participants reported whether the blue ball would reach the red or green semicircular arc (randomly oriented, equally dividing the disc edge). (B) Time-resolved decoding accuracy (multi-class SVM) of physical variables using MEG signals as predictors, based on all trials. Color codes represent physical variables: red for the blue ball’s initial motion direction γ, blue for the mass ratio k, and green for the relative angle α. The two continuous circular variables (γ and α) were evenly divided into four groups according to the angle values, while the mass ratio k were inherently segregated into three groups as dictated by the experimental design. Solid dark lines represent the physical simulation condition, while dashed light lines represent the background pattern condition. Dark and light symbols above the plots indicate the significance of the corresponding signals compared to the chance level, while black symbols denote significant differences between the physical simulation and background pattern conditions (n = 17 participants, light-shaded region indicates SEM, p < 0.05, cluster-based correction). (C) Decoding accuracy of the blue ball’s final motion direction γ′ using collision trials (because the final direction is defined only after a collision changes the ball’s trajectory). Similar to γ and α, γ′ were evenly divided into four groups. Same convention as (B), but this analysis used either MEG data or eye movement data (2-dimensional eye positions) as predictors. The arrows indicate the initial time points at which the decoding accuracy of γ′ shows significant differences between the physical simulation condition and the control condition (= 0.88 s for the MEG signal and 1.28 s for the eye movement signal). (D) Decoding accuracy of the participant’s final choice (binary classification SVM). Participants reported using their right index and middle fingers for red and green targets, respectively. Note that the red and green arcs in the visible and the invisible epochs are randomly rotated across trials.

To examine time-resolved decoding of variables under both conditions, we employed a multiclass support vector machine (SVM) classifier to decode each physical variable (Fig. 4B; detailed classification criteria in figure legend). Time-resolved decoding accuracy revealed distinct neural representations between physical simulation and control conditions. For the initial motion direction *γ* of the blue ball, decoding accuracy in the physical simulation condition exhibited a unimodal profile that peaked at 0.32 s (Fig. 4B, top, solid line; first significant time point = 0.12s; 17 participants, cluster-based correction, *p* < 0.05), followed by a gradual decline. In contrast, the background pattern condition showed a bimodal pattern: its first peak coincided with the physical simulation condition, but a second peak emerged after stimulus occlusion (Fig. 4B, top, dashed line; 17 participants, cluster-based correction, *p* < 0.05). This suggests that participants maintained a representation of initial direction during the invisible epoch in the control condition, as the information was required for task execution. Conversely, the absence of a second peak in physical simulation condition may reflect collision-induced directional updates, where the initial direction became behaviorally irrelevant after collision. Moreover, compared to the background pattern condition, both relative angle *α* and mass ratio *k* demonstrated significantly stronger decoding accuracy in the physical simulation condition, with prolonged temporal persistence (Fig. 4B, middle and bottom panels). Critically, given identical visual inputs across conditions, these MEG results—converging with the behavioral and fMRI findings—demonstrate that the brain dynamically represents relational physical variables only during physical simulation of collisions, which intrinsically requires tracking inter-object relations.

To test whether the predicted collision outcome—the blue ball’s final motion direction *γ′*—was represented specifically in the physical simulation condition, we used the *γ′* value from each physical simulation trial as the decoding target for the background trial with identical pre-occlusion stimuli. This means that the background-condition analysis in Fig. 4C used the matched collision-induced *γ′* as the decoding target, rather than the unchanged initial direction *γ* decoded in Fig. 4B. This matching was possible because the pre-occlusion stimuli were identical across conditions, allowing a one-to-one trial matching. Decoding results revealed significantly stronger accuracy in the physical simulation condition than in the background condition (Fig. 4C, top; 17 participants, cluster-based correction, p < 0.05, one-tailed Student’s t-test), with divergence emerging at ∼0.88 s.

We conducted analogous decoding analyses on eye-tracking data (Fig. 4C, bottom; 17 participants, cluster-based correction, *p* < 0.05), focusing on gaze coordinates (x, y). Decoding of blue ball’s final motion direction *γ′* revealed significant condition divergence at 1.28 s. Therefore, both neural and eye movement decoding results indicate that participants’ predictions about post-collision trajectories emerged at a relatively late time point. Notably, the average actual collision time across all physical scenes was 1.01 s, suggesting that participants’ mental simulation dynamics may closely align with the real-world physical dynamics—a correspondence that will be systematically examined in the next section. Importantly, the neural decoding results remained robust after regressing out the eye-movement signals from raw MEG data (Fig. S6C), suggesting that these effects were not primarily driven by oculomotor confounds.

### Dual-timescale dynamics of internal physical simulation

In our paradigm, the collision-induced directional change provides a temporal anchor point, allowing us to pinpoint when the brain updates its internal simulation of motion dynamics in response to collision events. In fact, under collision conditions, the emergence of the post-collision direction representation *γ′* exhibited a clear temporal delay relative to the onset of initial direction *γ* decoding (Fig. 4B, C).

To precisely quantify the collision-induced representational transition, we further employed ridge regression models to decode the initial *γ* and final *γ′* motion directions (see Methods). This continuous decoding approach (vs. categorical methods like SVM in Fig. 4) preserved fine-grained directional information, enabling precise detection of the exact moment of neural representational transition (Fig. 5A, bottom). The transition, as quantified by the crossover point between *γ* and *γ′* decoding error curves (1.10 s, 95% CI: 0.96–1.14 s), closely matched the average collision time of all physical scenes (1.01 s), suggesting temporal alignment between motion simulation updates and actual physical events. A parallel analysis of eye-tracking data revealed a similar transition pattern (Fig. S7A), consistent with prior evidence suggesting that eye movements can reflect internal simulation processes in the brain^33^. Importantly, the representational transition pattern was replicated in eye-movement**–**corrected MEG data (Fig. S6D), where horizontal and vertical eye positions were regressed out at the single-subject level prior to all analyses, supporting the conclusion that the observed transition was not primarily driven by oculomotor artifacts.

**Figure 5.**
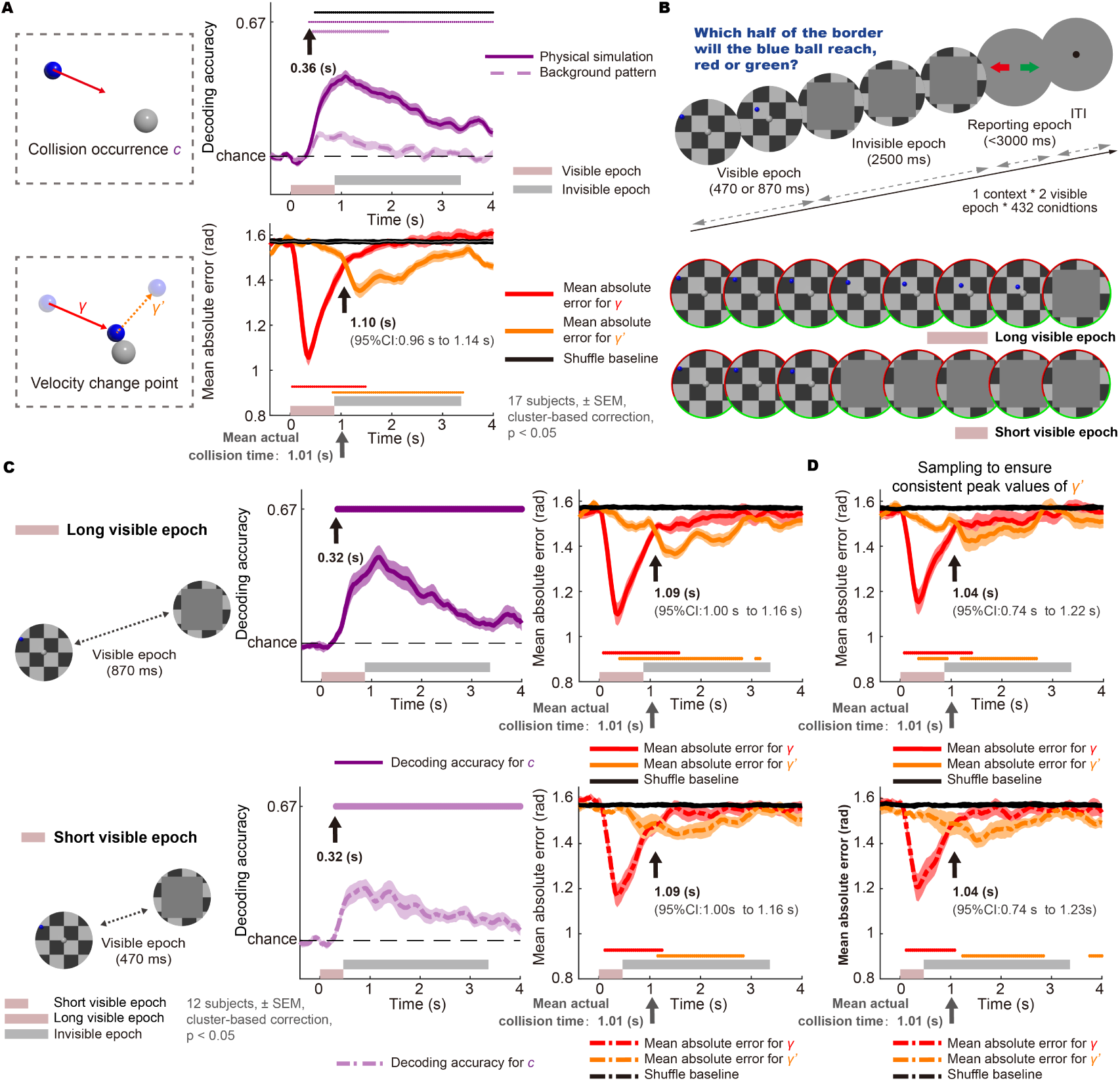
Internal physical simulation employs dual time scales. (A) Top: Decoding accuracy of collision occurrence using a binary SVM. A significant representation of collision occurrence in the physical simulation condition emerged at 0.36 s (indicated by the arrow). A significant divergence in collision representation between the two conditions emerged at 0.48 s (black symbols at the top). Bottom: Horizontal and vertical components of the blue ball’s motion direction were decoded using ridge regression. Decoding errors were quantified as mean absolute angular differences between predicted and actual directions. The decoding error curves for the motion direction of the blue ball before γ and after γ′ the collision intersect at 1.10 s (95% CI, bootstrapping; see Methods). The mean actual collision time across all stimuli is 1.01 s (SD=0.05). For both top and bottom: n = 17 participants; p < 0.05; light-shaded region indicates SEM; cluster-based correction applied. (B) Identical physical scenes were presented for different durations before occlusion. Both the long-stimulus condition (870 ms, identical to that of the main experiment) and the short-stimulus condition (470 ms, approximately half the duration) belonged to the physical simulation condition. No background pattern condition was included in this experiment. (C) Same as (A), but for the two stimulus conditions in (B) (n = 12 participants, p < 0.05, light-shaded region indicates SEM, cluster-based correction). (D) Decoding errors for the motion directions of the blue ball, using trials selected to match the distribution of minimum decoding errors (i.e., the trough of the decoding error curve) across the two stimulus conditions in (B).

The representational transition between *γ* and *γ′* at 1.10 s explicitly indicates that the brain encoded the trajectory dynamics of collision events. However, when directly decoding collision versus non-collision using SVM, we found that the neural representation of collision occurrence emerged at a remarkably earlier time point than the representational transition (0.36 s; Fig. 5A, top, solid line). This representation of collision occurrence cannot be attributed solely to low-level visual differences. Although some collision decoding persisted in the background pattern condition (Fig. 5A, top, dashed line), possibly reflecting subtle visual differences in the visible epoch between collision and non-collision trials, the substantially enhanced decoding accuracy in the physical simulation condition*—*despite identical visual stimuli across conditions*—*clearly demonstrates the engagement of internal simulation processes beyond basic visual processing.

The mismatch between early representation of collision occurrence and representational transition of *γ* and *γ′* hints at the possibility that the representational transition may actually occur well before the actual collision time, but remains masked by strong sensory inputs. In fact, the stimulus will not be occluded until 0.87 s, close to the mean collision time of 1.01 s. Thus, the apparent alignment between simulated and actual collisions might reflect the unmasking of pre-computed predictive signals following sensory withdrawal.

To test this possibility, we implemented a follow-up experiment that contrasted two conditions differing only in the duration of the visible epoch: a long visible epoch (0.87 s; same as in the main experiment) and a short visible epoch (0.47 s) (Fig. 5B). Notably, both conditions produced nearly identical neural dynamics: (1) the representation of collision occurrence emerged consistently at 0.32 s, and (2) the transition point in the decoding of the blue ball’s motion direction—reflecting the shift from pre-to post-collision representation—occurred at the same latency relative to stimulus onset (1.09 s in both conditions; 95% CI: 1.00–1.16 s), closely aligned with the actual collision time (Fig. 5C; see Fig. S7B for eye movement analyses). This duration-invariant temporal profile clearly argues against the unmasking account, which would otherwise predict a substantially earlier transition (predicted shift: *Δt* ≈ 0.4 s) under shorter stimulus exposure. The absence of such a shift indicates that the ∼1 s transition point reflects an internal simulation process rather than a sensory-driven unmasking effect.

While the transition timing from *γ* to *γ′* was consistent across long and short stimulus durations, we observed reduced peak decoding performance for post-collision motion direction *γ′* in the short-duration condition. The smaller amplitude of decoding accuracy may have delayed the intersection of the *γ* and *γ′* decoding curves in this condition. To isolate the amplitude effect, we selected a subset of trials where the distributions of *γ′* decoding error minima (i.e., troughs of decoding error curves) are comparable across conditions (see Methods). The preserved temporal alignment of collision-related transitions under this stringent control (Fig. 5D) rules out decoding amplitude artifacts.

Together, these findings demonstrate that during internal physical simulation, the brain operates on two distinct timescales: a real-time simulation that closely tracks the dynamics of the external physical world even during stimulus occlusion, and an early predictive signal that rapidly estimates high-level properties of the scene, such as collision occurrence.

### Predictive allocation of simulation resources

Notably, empirical evidence strongly supports a real-time simulation framework. Behavioral studies in humans and non-human primates, along with electrophysiological recordings in the monkey cortex^34,35^, demonstrate that the brain implements a form of dynamic inference or real-time simulation, continuously tracking object states rather than computing direct transformations from initial to final states. While our MEG results corroborate this real-time simulation mechanism, they further reveal that the prediction of collision occurrence emerges ∼700 ms before physical contact (Fig. 5). This temporal lead prompts a key question: what computational role does the early predictive signal serve?

Note that the brain’s internal simulations are inherently noisy—a fundamental constraint arising from spiking variability in neural circuits^36^. However, not all simulated physical events are equally susceptible to noise. Crucially, the difference between non-collision and collision represents a transition from a noise-insensitive to noise-sensitive regime: In collision trials, the blue ball’s final trajectory exhibits much higher sensitivity to perturbations in the initial motion direction (e.g. caused by noise) than in non-collision trials (Fig. S8, right column, showing a higher derivative of exit angle with respect to initial direction in collision trials). This dichotomy creates a computational demand: if neural systems can modulate noise levels (e.g., through attentional gain control^37^), prioritizing noise reduction during the early phase of physical simulations would disproportionately improve behavioral accuracy in collision versus non-collision trials. This prioritization mechanism could explain the early predictive signal observed.

To formalize this hypothesis, we developed computational models (Fig. 6A) grounded in the Intuitive Physics Engine (IPE) framework, which is essentially a probabilistic generative model that simulates physical processes through step-wise, procedural updates^24^. At each simulation step, multiple Monte Carlo samples are drawn to capture the stochasticity of internal predictions. Latent state features—such as object position and motion direction—are represented as probability distributions and updated according to physical rules. Within this framework, process noise is naturally incorporated into the probabilistic state representation: at each timestep, independent Gaussian noise with standard deviation *σ(t)* is added to each Monte Carlo sample of the latent features, reflecting the inherent uncertainty in internal physical simulations. Building upon psychophysical evidence that attentional resources regulate noise magnitudes in working memory^38–41^, we modeled noise controllability through dynamic resource allocation, whereby the available resources at time t, denoted *R(t)*, modulate the process noise *σ(t)* via a power-law relationship:

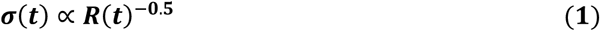

**Figure 6.**
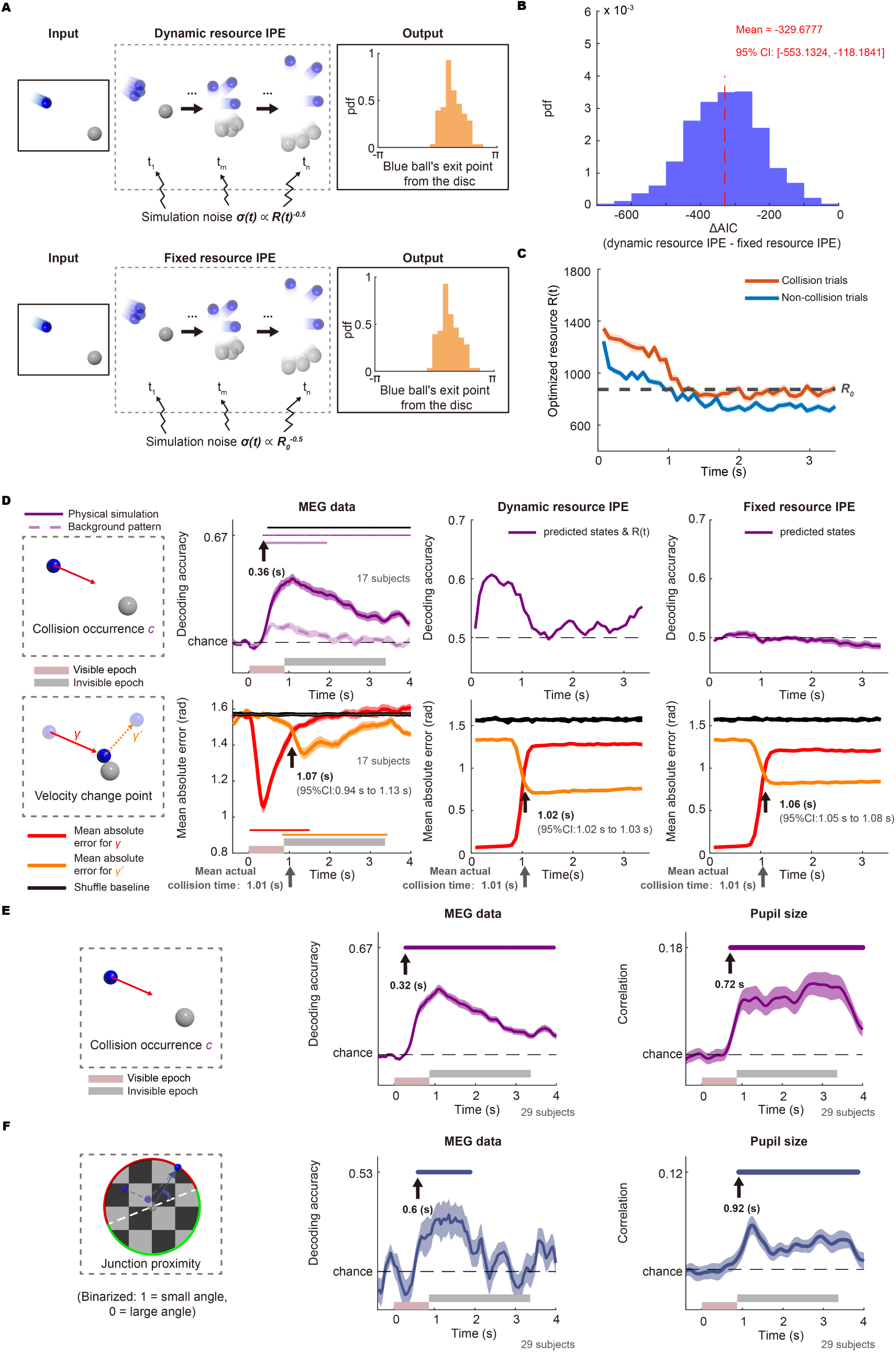
Emergence of dual time scales in dynamic resource IPE model. (A) Model design of dynamic and fixed resource IPE. (B) Distribution of ΔAIC (dynamic resource IPE – fixed resource IPE) obtained using the bootstrap method (1,000 iterations). Negative ΔAIC values indicate better performance of the dynamic resource IPE. The red dashed line indicates the mean ΔAIC across bootstrap iterations. (C) Time course of single-trial resource allocation estimates obtained by optimizing the dynamic resource IPE model with the best fitting parameters, averaged separately for collision and non-collision trials. Light-shaded region indicates SEM. A dashed horizontal line indicates the fitted constant resource level (R₀) from the fixed resource IPE model, providing a reference for comparison. (D) Same convention as Fig. 5A. Columns represent decoding results based on the following: 1) MEG neural activity (identical to Fig. 5A); 2) predicted states and R(t) from the dynamic resource IPE model; 3) predicted states from the fixed resource IPE model. (E) Left: Collision occurrence is coded as 1 for collision trials and 0 for non-collision trials. Middle: decoding accuracy of collision occurrence using a binary SVM. Significant difference from the chance level is indicated at the top (one-sample, one-tailed t-test, p<0.05). Right: Pearson correlation between pupil size and collision occurrence. Significant difference from 0 is indicated at the top (one-sample, one-tailed t-test, p<0.05). For middle and right panels: n = 29 participants (pooled from two MEG studies, all long-visible-epoch trials), p < 0.05, cluster-based correction. Analyses use physical simulation condition only. (F) Left: junction proximity coding based on exit angle relative to red/green junction: angles in the lower 50% of the sorted values coded as 1, others as 0. Middle: decoding accuracy of junction proximity using a binary SVM. Significant difference from the chance level is indicated at the top (one-sample, one-tailed t-test, p<0.05). Right: Pearson correlation between pupil size and junction proximity. Significant difference from 0 is indicated at the top (one-sample, one-tailed t-test, p<0.05). For middle and right panels: n = 29 participants, p < 0.05, cluster-based correction. Analyses use physical simulation condition only.

This implementation constitutes the dynamic resource IPE model. As a control, we introduced a fixed resource IPE model, where the resource maintains a constant level of *R_0_* throughout the simulation. In line with the standard IPE framework, both models output a probability distribution over the predicted exit point of the blue ball from the disc.

To model the brain’s dynamic resource allocation process, we developed a normative optimization framework that balances two competing demands—the cost of deploying attentional resources and the accuracy of simulation outcomes^42^. Critically, the resource allocation profile *R(t)* emerges as the solution to this optimization process rather than being externally specified. This approach builds upon van den Berg & Ma’s (2018) seminal work on set-size effects of working memory^43^, but crucially extents their single-trial resource allocation framework by introducing dynamic temporal allocation within trials.

The objective function of the dynamic resource IPE model is as follows:

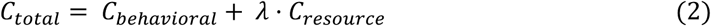

This total cost function *C_total_* reflects the trade-off between simulation accuracy (behavioral cost) and resource expenditure (resource cost), where the constituent terms are defined as follows:

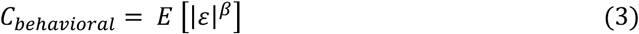

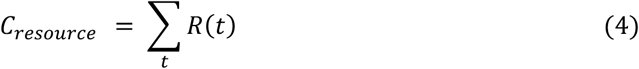

Here, *ε* indicates the difference between simulated and actual outcomes, *β* serves as an exponent transforming behavioral error into cost, and *λ* defines the relative importance of resource cost versus behavioral error.

The two free parameters *λ* and *β* were determined through Bayesian optimization^44^ to maximize the log-likelihood (LLH) of participants’ behavioral responses. Specifically, for each candidate (*λ*, *β*) pair, the model derives the optimal resource allocation *R(t)* that minimizes *C_total_*. Based on this optimized *R(t)*, the model generates predicted distributions of the blue ball’s exit angle. The log-likelihood of the observed behavioral data is then evaluated under these predictions. Parameters *λ* and *β* are iteratively updated to maximize the aggregate LLH across all trials and participants. In contrast to the dynamic resource IPE, the fixed resource IPE model contains only one free parameter: a time-invariant resource level *R_0_*, which is optimized using the identical Bayesian procedures.

We used the Akaike Information Criterion (AIC)^45^ and the Bayesian Information Criterion (BIC)^46^ to compare the performance of the two models. Bootstrap analysis consistently demonstrated the superior performance of the dynamic resource IPE over its fixed counterpart (Fig. 6B; Fig. S9A). In addition, cross-validation analysis yielded results consistent with both AIC and BIC (Fig. S9B), supporting the robustness of the model comparison. Furthermore, we compared the mean absolute error (MAE) of the two models against a baseline and the human behavioral noise floor (Fig. S9C). Both models performed significantly better than chance, yet a gap remained relative to the noise floor observed in human behavior.

For each time point, we computed the mean optimized resource (derived from our model with the best fitting parameters *λ* and *β*) by averaging across trials separately for collision and non-collision conditions (Fig. 6C). The fixed resource estimate, *R_0_*, is shown as a dashed line. Both collision and non-collision dynamic resource curves lie above *R_0_* during the early phase and fall below it after approximately 1 s, close to the actual collision time (1.01 s). Within this dynamic allocation, collision trials initially receive higher resources than non-collision trials, and this difference gradually diminishes over time. These results indicate that when resources can be dynamically allocated, the brain initially invests more—differentiating collision from non-collision conditions—and subsequently reduces allocation as the trial unfolds.

Our collision occurrence decoding analysis (Fig. 6D, top) revealed that only the dynamic resource IPE—incorporating both predicted states (velocity, mass etc.) and the time-varying resource *R(t)*—generated robust early representations of collision occurrence. The fixed resource IPE model’s weak collision representations mirrored the background condition result in MEG signals (dashed line in the 2^nd^ column of Fig. 6D), suggesting that this minimal decoding capability stems from incidental visual differences rather than true physical simulation.

To clarify the contributions of different model components, we performed ablation analyses (Fig. S10). When decoding from predicted states alone in the dynamic resource model (excluding *R(t)*), the early predictive signal was markedly reduced. Decoding from *R(t)* alone, or from predicted states in the fixed resource model augmented with *R(t)* from the dynamic resource model, largely recovered the robust early predictive performance observed in the full dynamic resource model. These results indicate that early predictive collision representations are driven primarily by the dynamic allocation of resources, captured by *R(t)*, rather than by the predicted physical states themselves. Crucially, *R(t)* in the dynamic resource model should not be interpreted as encoding explicit collision labels; rather, it reflects the model’s dynamically computed allocation of internal simulation resources under a resource–accuracy trade-off policy. This dynamic allocation provides a temporally adaptive, task-dependent dimension along which collision and non-collision trials can be distinguished early in the trajectory, thereby linking dynamic resource allocation to the rapid predictive collision signal.

For motion direction decoding (Fig. 6D, bottom), both models successfully captured the expected temporal dynamics: initial decoding of initial direction transitioning to final direction representation around 1s, consistent with the MEG findings. This demonstrates IPE framework’s inherent capacity for real-time motion simulation, independent of its resource allocation mechanism.

Notably, the gradual decline of optimized resources over time in the dynamic resource IPE model should not be interpreted as evidence that internal simulation after occlusion requires fewer attentional resources than perceptual processing before occlusion. Rather, this pattern partly reflects a simplification of this model: it relies on forward simulation from trial onset and does not explicitly distinguish between visible and invisible periods. In such a model, noise introduced earlier in the trajectory has a larger impact on the final simulation outcome than noise introduced later. To further account for the distinct visible and invisible phases of the experiment, we implemented an extended perceptual-simulation IPE model (Fig. S11). In this model, state estimates during the visible period are updated from the ground-truth physical state at each time step with perceptual noise modulated by resource allocation. Once occlusion begins, the model switches to forward simulation, with simulation noise also dependent on resources. This extension showed that resource allocation can be adjusted predictively before occlusion onset, while additional resources can be allocated during occlusion when perceptual information is no longer directly available, consistent with human attentional patterns observed in multiple-object tracking studies^47^.

Within our dynamic IPE framework, resource allocation is modeled as an unobserved latent variable. To ground it empirically, we examined pupil size—a well-established physiological marker of arousal that correlates with heart rate and skin conductance—as a proxy for attentional resource allocation^48–53^. Notably, pupil size showed significant positive correlations with collision occurrence starting at 0.72 s (Fig. 6E, Fig. S12), preceding by >300 ms the directional transition point observed in both MEG (1.10 s, Fig. 5A) and eye movement decoding analyses (1.13 s, Fig. S7A). This temporal precedence supports our model’s core proposition: predictive signals emerge early to guide dynamic resource allocation for real-time physical simulation.

Our computational modeling and pupil-size measurements support that the brain prioritizes simulation precision under conditions where small trajectory variations—caused by internal simulation noise—lead to disproportionately large outcome differences, as occurs in collision trials (Fig. S8). To test whether this principle may generalize beyond elastic collisions, we leveraged another aspect of our experimental design: the final response required participants to transform their predicted exit angle into a binary choice based on the red/green semicircular junction points. Crucially, when the predicted exit point approached the junction between color regions, the perceptual decision became more sensitive to subtle variations in the simulated trajectory, thereby demanding higher precision in physical simulation. Indeed, MEG patterns showed significant decodability of junction proximity, with pupil size positively correlating with proximity (Fig. 6F; see Fig. S13 for a breakdown into collision and non-collision trials). Moreover, this discrimination effect emerged later in time than collision-related signals (junction proximity vs. collision: 0.60 s vs. 0.32 s for MEG; 0.92 s vs. 0.72 s for pupil size), mirroring the physical event sequence (collision precedes disc-edge reaching). This temporal dissociation supports dynamic, cascaded allocation of cognitive resources during physical inference.

Consistent with this dynamic resource allocation account, fMRI revealed that the encoding of collision occurrence—and of the relational variables directly tied to collision occurrence—was primarily localized to higher-order cognitive regions (Fig.3D; Fig. S2B, C), which are well-positioned to coordinate cognitive resources. Moreover, in collision trials, relational variable representations were markedly stronger than in non-collision trials (Fig.3D), consistent with the idea that, under our proposed framework, collision conditions elicit greater cognitive resource allocation to these variables—especially the mass ratio *k*, which was identically distributed across collision and non-collision trials.

### A unified neural representation of simulated motion across collision and non-collision events

If the brain implements efficient physical inference through hierarchically organized predictive control that dynamically allocates cognitive resources, it would benefit from maintaining a unified representational format of motion-relevant physical variables, which provides a common basis for dynamic resource allocation and thereby enables rapid, accurate adaptation to diverse physical scenarios. This should be most evident during simulation, especially when visual input is occluded and internal dynamics dominate. To test this prediction, we conducted temporal generalization analyses to examine whether neural codes for object motion trained in one condition can generalize to another—for example, from collision to non-collision trials, or from the physical simulation condition to the background-pattern control. This approach allows us to assess whether the brain maintains a unified, simulation-driven code that supports flexible prediction across different physical scenarios.

We first analyzed non-collision trials from the physical simulation condition, which required maintaining *γ* representation without collision simulation throughout the invisible epoch. Temporal generalization decoding of the blue ball’s initial motion direction *γ* revealed two distinct decoding patterns in the temporal generalization matrix —an early and a late decoder (Fig. 7A, left column; 17 participants, cluster-based correction, p < 0.05)—that failed to generalize to each other’s time windows. The switch time point between the two decoders roughly aligned with the boundary between the visible and invisible epochs. This pattern was similarly observed in the background pattern condition (Fig. 7A, middle column), and even in cross-condition decoding between the two conditions (Fig. 7A, right column).

**Figure 7.**
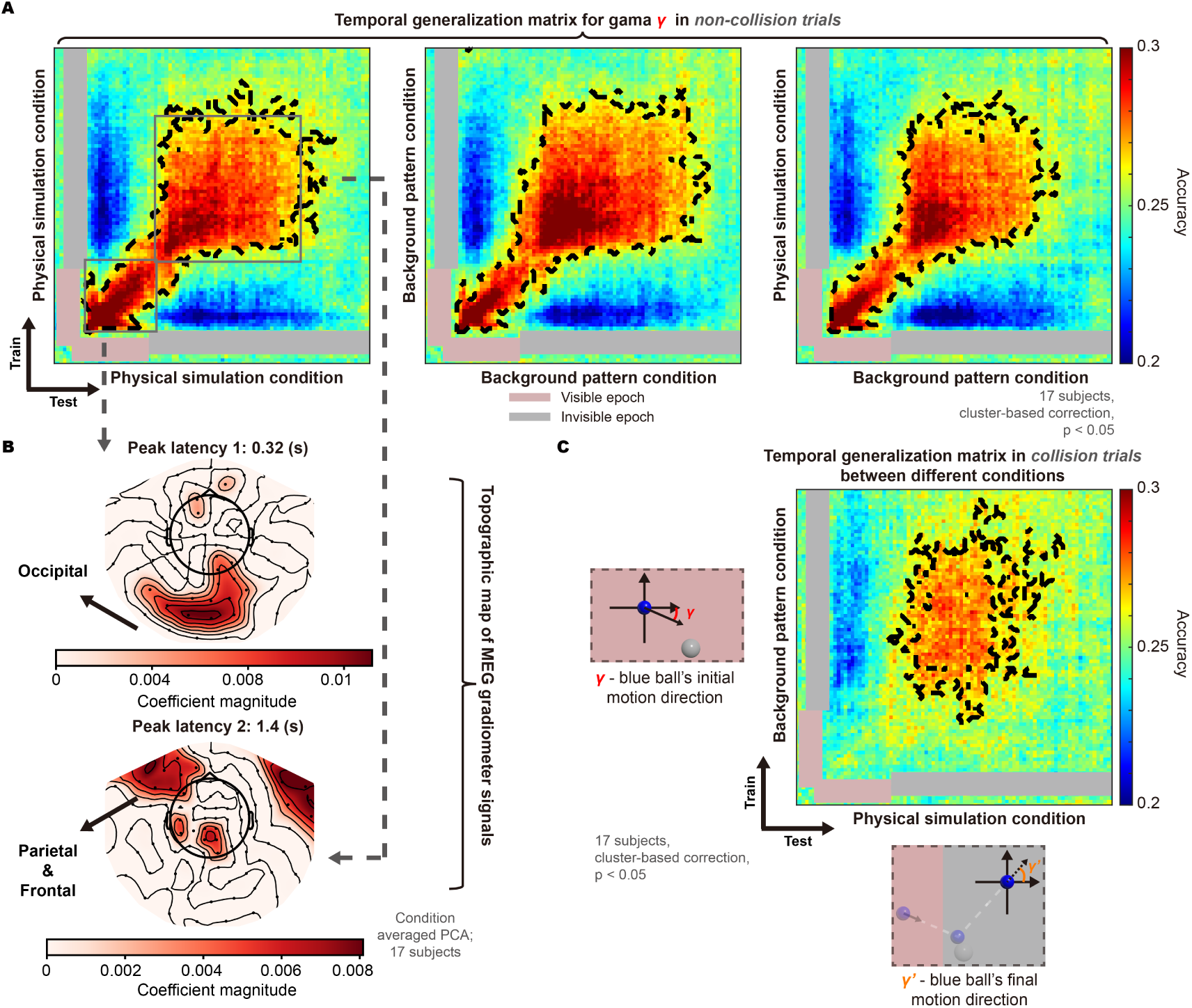
Temporal and condition generalization analysis. (A) Temporal generalization matrices quantifying the initial motion direction γ of the blue ball in non-collision trials. The left and middle panels depict within-condition generalization (left: physical stimulation; middle: background pattern), whereas the right panel demonstrates cross-condition generalization between the two experimental conditions. The black contours enclose temporal clusters with above-chance accuracy (n = 17 participants, p < 0.05, cluster-based correction). (B) Using MEG gradiometer data, a condition-averaged PCA method was applied to visualize the contribution of different spatial locations to encoding. The coefficient magnitude reflects the strength of neural involvement in encoding the variable at each location, with a 0.95 quantile threshold for selection. This analysis focused on the neural activity at the peak latencies of the two clusters in the temporal generalization matrix. Occipital regions are the predominant source at the earlier peak (= 0.32s), while parietal and frontal regions are the primary contributors at the later peak (= 1.4s). (C) Same convention as (A), but this analysis used collision trials from the physical simulation condition, along with matched trials from the background pattern condition that shared identical visual inputs. A classifier was trained to decode the blue ball’s initial motion direction γ in the background pattern condition, and tested on its final motion direction γ′ in the physical simulation condition.

To better understand the differences between the two decoders, we analyzed MEG gradiometer activity at each decoder’s peak decoding time point to identify their spatial patterns, as gradiometers offer greater sensitivity to local signal variations than magnetometers. Condition-averaged PCA revealed that *γ* encoding at 0.32 s (peak of early decoder) primarily involved posterior occipital cortex, while by 1.4 s (peak of late decoder), the representation had shifted anteriorly to frontal-parietal regions (Fig. 7B). This spatial redistribution aligns with a transition from sensory processing (early) to internal simulation (late), and is consistent with our fMRI finding that *γ* representation involved frontal-parietal regions in non-collision trials (Fig. 3D, bottom).

A critical remaining question is whether the brain utilizes a unified representation that generalizes between *γ* and *γ′* during motion simulation (the late decoder). To test this, we used the decoder trained on *γ* in non-collision trials to decode the simulated post-collision motion direction *γ′* in collision trials (both within the physical simulation condition). The results demonstrated successful generalization between the variables (Fig. S14).

To further rule out sensory confounds, we compared the collision trials in the physical simulation condition with perceptually matched but behaviorally distinct trials in the background pattern condition. Crucially, despite identical visual inputs, they differed in task requirements: the physical simulation condition required collision processing, whereas the background condition did not. A *γ*-direction decoder trained on background pattern trials also successfully generalized to *γ′* in the physical simulation condition (Fig. 7C; 17 participants, cluster-based correction, *p* < 0.05).

These results confirm our prediction: the brain indeed maintains a unified, simulation-driven representational format of object motion that generalizes across physical events and contexts (e.g., collision vs. non-collision; physical simulation vs. background pattern condition). This unified code provides the common basis for dynamic resource allocation, supporting efficient internal physical simulation and predictive control across diverse and changing environments.

## Discussions

In this paper, we combine a parameterized ball-collision paradigm with multimodal neuroimaging (fMRI/MEG) to elucidate how the human brain orchestrates physical simulations. Through systematic manipulation of physical variables, we reveal the following principles: 1) **Spatial segregation**. Physical variables exhibit a hierarchical organization, where object-specific features (*γ* and *γ′*) are encoded in sensorimotor cortices and relational variables (*α* and *k*) engage high-level cognitive regions. 2) **Temporal multiplexing.** Neural dynamics unfold at dual timescales: a slow simulation tracking real-time collision dynamics and a fast predictive signal (∼700 ms pre-collision) anticipating critical contacts. This architecture aligns with an extended IPE framework. In this framework, the brain dynamically regulates the allocation of resources to optimize the accuracy-cost tradeoff by prioritizing simulation precision based on sensitivity-critical variables (Fig. 8). Here, sensitivity-critical variables refer to parameters critically modulating system robustness, where minor state perturbations produce disproportionately large effects at specific critical values of these variables.

**Figure 8.**
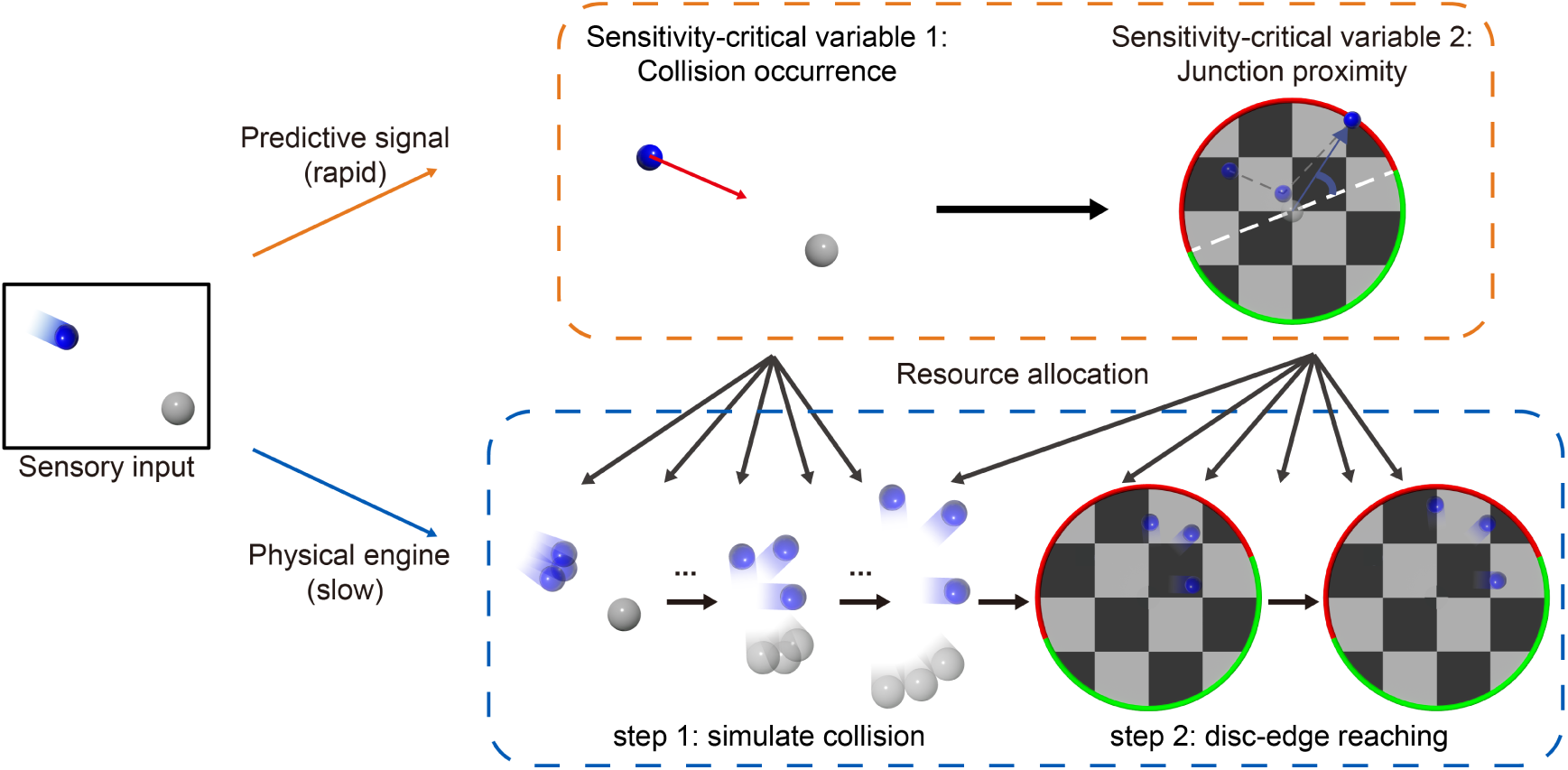
Predictive control dynamically regulates resource allocation in real-time physical simulation. The paradigm comprises two sequential stages: first, participants mentally simulate a collision and predict the post-collision motion trajectory of the blue ball; second, they convert the predicted exit angle of the blue ball into a red/green choice by comparing it to the disc’s red-green boundary position. This task engages a real-time internal physical engine in the brain, which simulates and predicts object dynamics with temporal characteristics closely aligned with real-world physical processes. In parallel, a faster predictive component continuously estimates sensitivity-critical variables essential for the system’s robustness. We propose that such predictive representations dynamically regulate the allocation of cognitive resources available to the real-time physical engine.

The ability to flexibly predict and infer object movements represents a fundamental survival skill, observed across species from non-human primates to human infants^2–6,33–35^. A central cognitive theory posits that these computations rely on mental simulations mediated by a frontoparietal network—a distributed neural system that encodes physical variables and supports dynamic prediction^7,15–17^. However, this framework has traditionally treated the frontoparietal network as a unitary computational module, overlooking its intricate functional subdivisions and temporal dynamics during physical reasoning tasks.

Using physical variables as computational anchors, our findings reveal a spatiotemporally organized architecture of internal physical simulation. Although acquired independently, the fMRI and MEG data converge within a unified framework: rapid collision detection signals (MEG) selectively engage high-level cognitive regions (fMRI), which in turn orchestrate resource allocation for slower, precision-sensitive real-time simulation (modeling + pupillometry). Specifically, MEG revealed an early predictive signal encoding collision occurrence, temporally preceding the actual collision by 700 ms (Fig. 5A). In parallel, fMRI identified that regions involved in encoding collision occurrence largely overlapped with the executive control network (Fig. S4B, C), a core neural substrate of cognitive control^54,55^. This overlap suggests the predictive signals in physical inference are embedded within the brain’s broader cognitive control architecture, enabling flexible regulation of resources for simulation. These converging findings support the Dynamic Resource IPE model, which posits that the brain selectively allocates cognitive resources via top-down modulations to optimize the accuracy-cost tradeoff (Fig. 6).

Guided by the Dynamic Resource IPE framework, we uncovered a temporally cascaded resource allocation pattern. Convergent MEG and pupillometric evidence supported that, 200-300 ms after the initial collision detection signal, a secondary allocation phase emerged, dynamically regulated by the spatial proximity between the final exit point and the red/green decision boundary (Fig. 6F, Fig. 8). Crucially, this sequential pattern nicely mirrors the physical event chronology, suggesting that even the rapid, resource-orchestrating predictive process embodies the causal structure of Newtonian dynamics. It is to our surprise that such intricate structures emerge out of a seemingly simple paradigm, implicating that ‘physical inference’ could constitute a naturalistic approach for examining complex, hierarchical cognitive architectures^56–58^.

Our dynamic resource IPE model optimizes a resource allocation profile *R(t)* by evaluating prediction accuracy against the known outcome. This use of the ground-truth outcome is a computational device for estimating an advantageous resource-allocation profile, and should not be taken to imply that human observers have access to the true outcome on the current trial. Rather, the optimization can be interpreted as a normative approximation to a resource-allocation policy acquired through long-term experience. Human observers do not have access to trial-specific outcomes, but extensive exposure to physical events—such as object collisions or motion continuity in everyday life—enables the brain to learn probabilistic models that guide predictive simulation. In everyday experience, observers often encounter both the initial conditions and the subsequent outcomes of physical events, providing prediction-error feedback that can support learning of physical dynamics and of when additional computational resources are most useful. The observed early predictive signals and differential resource allocation in our experiment likely reflect these experience-driven strategies, rather than trial-specific knowledge, consistent with evidence from intuitive physics and predictive processing frameworks^24,59–62^. Thus, although our model is not intended as a full online learning mechanism, it provides a normative computational account of how learned physical expectations could guide dynamic allocation of cognitive resources during physical prediction.

In this study, our multimodal approach revealed a consistent dual-timescale pattern across both neural and behavioral dimensions. While MEG captured the temporal dynamics of neural processing, complementary behavioral measures—including eye position tracking and pupillometry—provided converging evidence. Notably, pupil diameter variations exhibited rapid predictive encoding of collision occurrence (Fig. 6E), paralleling the fast anticipatory components in MEG data. On the other hand, eye movement patterns, mirroring the slower MEG components, continuously tracked the ball’s simulated trajectory in real time (Fig. S7B). This cross-modal replication of the temporal dissociation robustly validates our hierarchical processing framework.

The observed 200-300 ms lag between MEG signals and pupil responses—present for both collision occurrence and junction proximity representations (Fig. 6E, F)— likely reflects hierarchical processing in physiological arousal modulation. Early MEG components encode sensory predictions, whereas delayed pupil dynamics index neuromodulatory mobilization involving noradrenergic and cholinergic pathways. The locus coeruleus (LC) acts as a critical interface mediating the translation of cortical predictions into autonomic responses. This temporal gap may correspond to: 1) Cortical-subcortical transmission time (frontal-LC pathway); 2) Neuromodulatory integration (LC’s noradrenergic release to pupil control circuits). Notably, while the observed delay magnitude aligns with prior evidence showing ≈300 ms latency between LC activity and pupil dilation under passive sensory paradigms^63^, the complete cortical-LC-pupil pathway timing during active physical inference requires further characterization.

The core contribution of our work lies in demonstrating that physical simulation in the brain operates as a multi-scale process: a slow simulation tracking real-time collision dynamics and fast predictive signals anticipating sensitivity-critical variables, such as collision occurrence. These predictive signals guide the dynamic allocation of cognitive resources to fine-tune real-time simulation fidelity. While real-time physical simulation is supported by prior empirical and computational studies^34,35^, its computational rationale appears paradoxical: Why simulate the physical scenarios in real time, even when they’re governed by familiar physical principles, rather than to precompute the outcome for faster responses?

The answer may lie in real-time simulation’s unique advantages. Unlike precomputed solutions limited to specific scenarios, real-time simulation enables dynamical tracking of evolving physical states and immediate adaptation to novel conditions. This is particularly beneficial in complex, nonlinear environments where small variations can result in disproportionately large differences in outcomes. This flexibility is further enhanced by dynamic resource allocation: by rapidly predicting sensitivity-critical variables, the brain strategically focuses cognitive resources where most needed, achieving an optimal efficiency-precision balance in physical reasoning.

Interestingly, this hierarchical progression—starting with high-level predictive signals about sensitivity-critical variables and leading to the refinement of lower-level simulations—echoes a principle reminiscent of the Reverse Hierarchy Theory (RHT) in visual perception^64,65^. In RHT, high-level representations guide the optimization of lower-level feature processing. Our findings extend this principle beyond visual perception to encompass internal physical simulation and provide empirical support for such top-down modulation in a complex, dynamic task.

This neurocomputational principle offers valuable insights for the development of more robust and adaptive AI systems. Traditional physics engines or predictive models in artificial agents typically rely on fixed-resolution simulations or end-to-end learning pipelines that inherently lack dynamic flexibility; however, like the human brain, AI systems also operate under resource constraints, highlighting the need for mechanisms that adaptively allocate computational resources. By incorporating a brain-inspired mechanism—where computational precision is continuously modulated according to the anticipated sensitivity of the system to perturbations—we could significantly enhance both the efficiency and generalization capability of AI agents in complex environments. For example, implementing sensitivity-based predictive control would allow artificial systems to intelligently allocate computational resources, concentrating processing power on precisely those critical moments or variables where prediction errors have the greatest impact. This mechanism could improve performance across various domains^66^, including embodied AI (particularly in physical embodiment scenarios), autonomous navigation, physical reasoning, and intuitive decision-making—especially under conditions of uncertainty or limited computational budgets.

## Methods

We conducted four complementary studies: a behavioral experiment (Study 1, *n* = 16; Fig. 2), an fMRI experiment (Study 2, *n* = 11; Fig. 3) and two MEG experiments (Study 3, *n* = 17; Figs. 4, 5 and 7, and Study 4, *n* = 12; Fig. 5). All studies employed modified versions of the same core experimental paradigm where participants observed videos of parameterized ball-collision scenarios. On each trial, an initially moving blue ball approached a stationary white ball, and participants predicted the subsequent trajectory of the blue ball.

### Participants

Sixteen healthy participants (eight males, eight females; mean age = 25.06 years, range = 22–30 years) completed the behavioral experiment. Eleven participants (three males, eight females; mean age = 25.36 years, range = 23–29) participated the fMRI experiment. Seventeen participants (eight males, nine females; mean age = 24.65 years, range = 23–31) and twelve participants (four males, eight females; mean age = 24.92 years, range = 24–28) participated in the first and second MEG experiments respectively. All experimental protocols were approved by the Biomedical Research Ethics Committee of the Institute of Neuroscience, Chinese Academy of Sciences. All participants had given written consent to the procedure in accordance with institutional guidelines and the Declaration of Helsinki. Participants were compensated at a rate of 1 Chinese yuan/minute for behavioral experiments and 2 Chinese yuan/minute for MEG/fMRI experiments. Sex/gender was not a factor in the study design and was not analyzed.

### Stimulus Design

All visual stimuli were constructed using Autodesk 3ds Max 2023 (Autodesk Inc., San Rafael, CA, USA), yielding 432 unique ball-collision scenes. Each video depicted an elastic collision between two balls on a frictionless platform, with stimuli subtending 14.5 × 14.5 degrees of visual angle.

Each video began with a white ball stationary at the center of a disc. A blue ball appeared at one of 12 equidistant positions (30 ° angular intervals) along the disc’s perimeter and moved at a fixed speed in a randomized direction. At a predetermined time point (0.87s after stimulus onset, termed the ‘visible epoch’), prior to the anticipated collision, the two balls were occluded (termed the ‘invisible epoch’, duration=2.5s). During this invisible epoch, participants mentally simulated the unfolding physical process to predict the outcome. In all experiments, participants were required to predict final exit point of the blue ball from the disc. Reporting procedures differed between behavioral and neuroimaging experiments.

To ensure a 50% collision probability across all experimental conditions, we parameterized the blue ball’s initial motion relative to the white ball within a coordinate system centered on the white ball. The relative angle *α*, defined as the angular deviation between the blue ball’s velocity vector and the inter-ball center vector, constituted a critical parameter governing whether a collision would occur. This angle was sampled from a Gaussian distribution calibrated to yield collisions in 50% of the trials. We further manipulated the mass ratio between the two balls (*k*=m_blue/m_white) through three experimental conditions: (1) *k*=3.375, (2) *k*=1, and (3) *k*=0.422. Under the assumption of identical material density between balls, these ratios were implemented by proportionally scaling the size of the white ball while keeping the blue ball’s physical dimensions unchanged throughout all trials.

Two experimental conditions were implemented: (1) the physics simulation condition and (2) the background pattern control condition. In the physics simulation condition, the white ball functioned as a physical entity possessing mass, where elastic collisions with the blue ball induced trajectory modifications governed by Newtonian mechanics. In contrast, in the background pattern condition, the white ball served merely as a visual element embedded within the static checkerboard background, thus producing no kinematic effects on the blue ball’s movement.

The behavioral and fMRI experiments employed only the physics simulation condition, whereas the MEG experiments incorporated both the physics simulation and background pattern conditions. Importantly, both conditions used identical initial parameters and contained 432 distinct videos each. As occlusion occurred before any ball-to-ball contact, the visual input during the visible epoch remained indistinguishable between conditions. Participants were informed of the experimental condition via a block-wise textual cue, allowing them to construct appropriate mental simulations: predicting either Newtonian collisions (physical simulation condition) or uniform linear motion (background pattern condition) during the invisible epoch.

### Pretraining

The pre-experimental protocol was implemented consistently across all four experiments. First, participants observed a one-minute, unoccluded demonstration of ball collisions to familiarize themselves with the physical setup of the scenario. They then performed practice trials identical to the formal experimental trials to acclimate to the occlusion paradigm.

### Study 1: behavioral experiment

#### Experimental procedures

In the behavioral experiment, participants were required to report the predicted exit point of the blue ball. The perimeter of the background disc was uniformly set to either fully red (visible + invisible epoch) or green (reporting epoch). When the invisible epoch ended, signaled by the perimeter’s color transition from red to green, participants clicked on the disc’s perimeter. The selected position was converted into an angle value in the world coordinate system centered at the disc’s origin.

#### Full Newtonian mechanics and perturbed models

To establish the ground truth, the exit point of the blue ball after collision was derived using momentum and kinetic energy conservation within the Newtonian mechanics framework for perfectly elastic collisions. The derivation yielded the exit angle as a function of *γ*, *α*, and *k* (*γ*: initial motion direction, *α*: relative angle, *k*: mass ratio). Three perturbed models were implemented: 1. Constant mass ratio: fixed *k* across trials (three distinct ratios generating three sub-models); 2. Random *α:* Trial-wise shuffling of relative angles; 3. Ignore white ball: Setting collision momentum transfer to zero.

### Study 2: fMRI experiment

#### Experimental procedures

The fMRI experiment employed an event-related design with jittered inter-trial intervals (ITIs) sampled from a uniform distribution ranging from 4 to 8 seconds (mean = 6 s) to minimize temporal expectation effects^67^. To dissociate motor-related signals from the cognitive processes involved in physics simulation, the behavioral reporting protocol was modified. Instead of reporting the precise exit angles, participants indicated whether the blue ball would reach the red or green semicircular border segment (randomly rotated on each trial) using right-hand button presses (index finger for red, middle finger for green). This design ensured that motor responses were color-contingent, while exit-angle predictions remained invariant to rotation, thereby effectively segregating simulation-related neural activity from motor execution signals.

## Data acquisition

All participants were scanned using a standard 32-channel phased-array head coil on a Siemens Tim Trio 3.0 T scanner (Erlangen, Germany). Whole-brain fMRI data were collected using a gradient-echo echo-planar imaging (EPI) sequence (TR = 2000 ms, TE = 30 ms, flip angle = 90°, spatial resolution = 3mm^3^, number of slices = 50, multiband factor = 2, phase encoding direction: anterior-to-posterior, echo spacing = 0.52 ms). A pair of gradient echo images (echo times: 8 ms and 10.46 ms) with the same orientation and resolution as the EPI images were acquired to generate a field map for distortion correction of the EPI images. High-resolution T1-weighted anatomical images were obtained using a MPRAGE sequence (TR = 2300 ms, TE = 3 ms, inversion time = 1000 ms, flip angle = 9°, spatial resolution = 1mm^3^, 176 sagittal slices). Each fMRI experiment lasted 90-120 minutes. Visual stimuli were presented using Psychtoolbox-3 (version 3.0.19.15) on MATLAB 2020b. Participants were allowed to move their eyes freely during the experiment, and no eye tracking was performed during the fMRI sessions. Eye movements were continuously recorded in the MEG experiment (see MEG Methods for details).

### Preprocessing

fMRI data were preprocessed and analyzed using SPM12 (Wellcome Department of Cognitive Neurology, London, UK). The first five volume images (5 TR, 10 seconds) of each run were discarded to allow for T₁ saturation and establishment of steady-state magnetization. All functional images were then realigned to the first volume to correct for head motion during scanning. Subsequently, volumes were undistorted based on the acquired field map. Finally, each participant’s T1-weighted anatomical image was coregistered to their mean functional image.

### Single-trial beta estimation

A general linear model (GLM) was applied to the preprocessed functional images using SPM. Separate regressors modeled each trial to estimate single-trial responses. Six motion parameters were included as nuisance covariates. The GLM design matrix was applied independently to each run and was convolved with a canonical hemodynamic response function (HRF) to model the BOLD response.

### Transformation from voxel-based to surface-based beta series

The estimated beta series were projected from volumetric space onto individual cortical surface meshes (vertex-based representation) using the participant-specific surfaces derived from their T1-weighted anatomical images^68^. These individual surface maps were then registered to a standardized template surface (e.g., fsaverage) and spatially smoothed with a 6 mm full-width at half-maximum (FWHM) Gaussian kernel via the CAT12 toolbox^69^. This process yielded standardized surface-based beta series for each participant, where trial-specific response magnitudes were represented by beta values at each vertex. These surface-based beta estimates were then used for subsequent univariate analyses and multivariate decoding.

### Whole-brain univariate ANOVA

Among the four physical variables, the blue ball’s initial motion direction (*γ*), relative angle (*α*), and post-collision direction of the blue ball (*γ′*) are continuous circular variables. Given their circular nature, conventional linear assumptions were inappropriate. We therefore adopted a whole-brain univariate ANOVA approach to identify brain regions exhibiting selective responses to each variable.

For each variable, trials were binned according to parametric variable. This approach capitalizes on the principle that vertices encoding a specific variable should demonstrate greater between-group response variance (across distinct values) than within-group variance (among similar values). Accordingly, we computed vertex-wise F-scores from the beta series to quantify this relationship.

The specific procedure involved Z-score normalization of individual surface-based beta series to account for response magnitude differences across participants, followed by cross-participant averaging of normalized beta series at each vertex location^70^. F-scores were then computed from the group-averaged beta series for each vertex, generating whole-brain F-statistic maps. Finally, false discovery rate (FDR) correction for multiple comparisons (p<0.05) was applied using MATLAB’s implementation.

### Whole-brain multivariate decoding

We performed whole-brain multivariate pattern analysis (MVPA) to decode the blue ball’s initial motion direction (*γ*). Two decoding schemes were implemented using linear support vector machine (SVM) classifiers: cross-trial-type decoding and within-trial-type decoding. Chance level was 25% for the four-class classification.

For cross-trial-type decoding, classifiers were trained on collision trials and tested on non-collision trials, and vice versa.

For within-trial-type decoding, classification was performed separately within collision and non-collision trials using a five-fold cross-validation procedure. Decoding accuracy was averaged across folds.

### Multivariate decoding of relative angle in non-collision trials

To assess whether relative angle (*α*) is represented in non-collision trials despite the absence of significant univariate tuning, we performed additional MVPA analyses to decode *α* specifically within non-collision trials. Decoding was conducted under two feature selection schemes. In the first scheme, the analysis was restricted to vertices that were significantly tuned to α in collision trials, as identified using a univariate ANOVA (α-selective vertices). In the second scheme, decoding was performed using all vertices across the whole brain. Classification was implemented using the same linear SVM procedure and five-fold cross-validation scheme as used for within-trial-type decoding above. Chance level was 25% for the four-class classification.

### Permutation-based test of spatial overlap across physical variables

To assess spatial overlap among cortical representations of different physical variables during collision trials, we constructed a null distribution using a permutation procedure. The spatial locations of vertices significantly tuned to each variable were randomly shuffled across the cortex while preserving the number of selective vertices for each variable, and this procedure was repeated 1,000 times. For each permutation, we counted the number of vertices selective for two or more physical variables, yielding a null distribution of multi-variable overlap the null hypothesis of random spatial arrangement. The observed overlap was then compared against this null distribution to assess statistical significance.

### MT ROI-based analysis

The MT ROI (middle temporal area, a motion-sensitive visual region) was defined using the Human Connectome Project (HCP) multi-modal parcellation (version 1.0). For each vertex, beta values were first z-scored across trials within individual participants, then averaged across all vertices within the MT ROI (separately for the right and left hemispheres), and finally averaged across participants to generate group-level responses.

### Overlap with naturalistic movie-derived functional parcellation

To quantify the hierarchical organization of neural representations, we aligned the cortical map of each physical variable with Rajimehr et al.’s (2024) 24-network parcellation derived from naturalistic movie fMRI. Both datasets were registered to the Freesurfer “fsaverage” template (32k vertices). Spatial overlap between brain areas showing significant tuning in whole-brain univariate ANOVA and the 24 functional networks were quantified using the Jaccard index (JI), defined as the ratio of the number of vertices in the intersection to that in the union between two vertex sets:

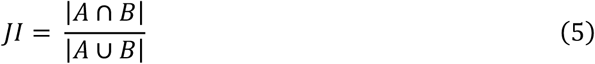

where A is the set of vertices significantly representing a given physical variable, and B is the set of vertices belonging to a given functional network.

### Study 3/4: MEG experiment

#### Experimental procedures

The MEG experiment employed a dual-condition design (physics simulation vs. background pattern). All participants completed the same 432 trials, randomly distributed across 12 runs with individualized trial sequences. Condition blocks were interleaved within runs and cued by textual prompts instructing participants to simulate either Newtonian collisions (physics condition) or uniform linear motion (background condition).

The MEG experiment adopted the same behavioral reporting protocol used in the fMRI experiment to dissociate motor-related signals from physics simulation processes. Instead of reporting the precise exit angles, participants indicated whether the blue ball would reach the red or green semicircular border segment (randomly rotated per trial) via right-hand button presses (index finger for red, middle finger for green).This design ensured that motor responses were color-contingent while exit-angle predictions were rotation-invariant, effectively segregating simulation-related activity from motor execution signals.

#### Data acquisition

Participants performed the tasks while seated inside an electromagnetically shielded room. The magnetic component of their brain activity was recorded using the 306-channel TRIUX^neo^ MEG system (MEGIN, Finland), which consists of 102 sensor triplets, each comprising one magnetometer and two orthogonal planar gradiometers. The brain signals were acquired at a sampling rate of 1000 Hz with hardware highpass filtering (0.1-330 Hz). Offline processing included head movement compensation via MaxFilter software (MEGIN, Finland) and interference suppression using temporal signal space separation (tSSS).

Continuous physiological monitoring encompassed electro-oculograms (EOGs) for oculomotor activity and electrocardiograms (ECGs) for cardiac rhythm recording. During the MEG acquisition, participants’ eye movements were not physically constrained, while EyeLink 1000 Plus system (SR Research, Canada) tracked comprehensive oculometrics including gaze position coordinates and pupil size dynamics over time.

#### Preprocessing

The remaining preprocessing steps were performed using the MNE-Python package^71,72^. Bad channels were identified and interpolated. Signal filtering included a high-pass filter (0.1 Hz cutoff), a low-pass filter (150 Hz cutoff), and a notch filter at 50 Hz and its harmonics (100, 150, 200, and 250 Hz) to eliminate power line noise. The data were decimated by a factor of 2, reducing the sampling rate from 1000 Hz to 500 Hz.

Independent Component Analysis (ICA) was applied to ocular (EOG) and cardiac (ECG) artifacts. Artifact-related components were identified using correlation-based methods implemented in MNE (i.e., find_bads_eog and find_bads_ecg), and excluded from subsequent analysis. Baseline correction was performed using the-0.2 s to 0 s interval. Finally, the data were further downsampled to 40 Hz and temporally smoothed using MATLAB’s smooth function (40 ms window radius) before decoding analyses.

#### Quantification of gaze tracking error

To assess potential differences in eye movements across trial types, we analyzed eye-tracking data recorded during the MEG experiment. For each participant, we computed a time-resolved tracking error defined as the Euclidean distance, in degrees of visual angle, between the participant’s gaze position and the instantaneous center of the blue ball on the screen. This analysis was restricted to time points during the visible epoch, as the target object was occluded during the invisible epoch. Tracking error was computed separately for each trial and then averaged as a function of time for collision trials and non-collision trials. This allowed us to quantify whether gaze behavior systematically tracked the blue ball and whether such tracking differed between collision and non-collision trials.

#### MEG residuals after eye-movement regression

Since natural viewing behavior was permitted during the experiment, we employed linear regression covarying out potential artifacts associated with eye movements. Eye-tracking signals (gaze coordinates on screen), which underwent identical smoothing and downsampling procedures as the MEG recordings, served as regressors to model MEG signal variance. All time-series data were z-scored prior to regression model fitting. The residuals from the regression were then used for further analysis.

#### Multi-class SVM decoding

The following variables were decoded using multi-class support vector machine (SVM) with one-vs-rest classification strategy: the blue ball’s initial motion direction (*γ*, 4 levels), the blue ball’s final motion direction (*γ′*, 4 levels), relative angle (*α*, 4 levels), mass ratio (*k*, 3 levels), motor response selection (*m*, 2 levels), and collision occurrence (*c*, 2 levels). Prior to model training, the data at each time-point were z-scored across trials. We implemented a cross-validation procedure with an 80:20% training-to-test split, repeated 100 times with random sampling. Decoding accuracy was temporally smoothed using a moving average with a 120 ms window radius and then averaged across participants. Statistical significance was determined using a cluster-based permutation test (p< 0.05). While γ′ decoding was performed exclusively on collision trials, all other physical variables were decoded using the complete trial set.

#### Temporal generalization analysis

The decoding function for the temporal generalization matrix employed a multi-class SVM approach (one-vs-rest classification), with cross-validation conducted using 80% of the data for training and 20% for testing, repeated for 100 iterations. Prior to model training, data at each time-point were z-scored across trials. The critical distinction resided in training/testing set selection: decoding was performed either (a) across distinct time points within identical conditions, or (b) across different conditions and time points. Statistical significance was determined using a cluster-based permutation test (p <0.05).

#### Topographical encoding of physical variables

To characterize spatial encoding patterns, we performed condition-averaged PCA on sensor-level data. The analysis utilized MEG gradiometer residuals (204 sensors) with eye movement artifacts regressed out using eye-tracking data as covariates. Gradiometers were selected for spatial pattern analysis due to their greater sensitivity to local cortical sources compared to magnetometers. The MEG residuals at each time point were z-scored across trials, and then averaged within experimental conditions grouped by the target physical variable (e.g. the blue ball’s initial motion direction). The resulting N × C matrix (conditions × channels) underwent PCA, where component loadings revealed spatial encoding patterns. Channels exceeding the 95th percentile of absolute loadings for the first principal component (PC1) were visualized.

#### Ridge regression

The internal simulated dynamics of the blue ball were modeled using ridge regression. This approach was applied to decode both the initial (*γ*) and final (*γ′*) motion directions of the blue ball. Two separate ridge regression models were trained to predict the horizontal (cosine) and vertical (sine) components of each angular variable. The regularization parameter *λ* was optimized through a logarithmic grid search spanning 100 values from10⁻⁴ to 10⁴. Cross-validation was conducted using 80% of the data for training and 20% for testing, repeated for 100 iterations. The predicted horizontal and vertical components were then converted to angular values (range:-π to π). Decoding performance was quantified using the mean absolute error (MAE) of the predicted angles, with statistical significance determined through permutation testing (1,000 label-shuffled iterations).

The intersection point of the decoding curves for initial (*γ*) and final (*γ′*) motion directions marks the time at which the brain transitions between representing the blue ball’s pre-and post-collision trajectories. The confidence interval of this intersection was estimated through a bootstrapping (1,000 iterations).

To minimize mutual predictability between the initial (γ) and final (γ′) motion directions, the decoding curves (Study 3; Fig. 5A, bottom) were generated using a subset of collision trials selected by a threshold criterion for absolute direction change. Specifically, we included trials where the blue ball’s direction change before versus after collision exceeded 0.3 radians (139 of 211 collision trials), with angular similarity decreasing from 0.48 before selection to 0.23 after selection. For comparison, the same analysis was replicated using all available collision trials (211 trials) following the procedure in Fig. 5A. Crucially, the intersection points of the decoding curves for initial (γ) and final (γ′) motion directions remained nearly identical (Fig. S7A), confirming the robustness of intersection point estimation to trial selection criteria.

To control for differences in peak decoding performance (minima of decoding error) of the blue ball’s final motion direction *γ′* between the long and short visible epoch conditions (Study 4; Fig. 5D), we matched trials by aligning their MAE distributions at the peak time point. Specifically, we binned the γ′ decoding error minima distributions for both conditions. For each bin, we randomly subsampled trials without replacement from the condition with more trials to match the count in the condition with fewer trials. This procedure generated matched distributions across conditions.

### Dynamic resource IPE model

#### Overview

The dynamic resource IPE model extends the IPE (Intuitive Physics Engine) framework by incorporating and dynamically regulating internal simulation noise. The IPE is formulated as a probabilistic generative model that simulates physical processes through step-wise procedural updates. At each timestep, latent state features—such as object position and motion direction—are represented as probability distributions and updated according to physical rules. To capture stochasticity of internal simulations, multiple Monte Carlo samples of the latent state are drawn at each timestep. Each sample evolves deterministically under the physical rules, but independent Gaussian process noise with standard devidation *σ_t_* is added to each sample, producing an ensemble of samples in latent feature space whose distribution reflects uncertainty in internal simulation. In our implementation, the model outputs a probability distribution over the exit angle of the blue ball.

For simplicity, the dynamic resource model assumes access to the true initial state of the scene without sensory noise. Uncertainty arises from inherent biological noise during mental simulation, reflecting intrinsic stochasticity in neural computations. This noise is modeled as a zero-mean Gaussian distribution N (0, *σ(t)*^2^), where the standard deviation *σ(t)* is dynamically regulated by the computational resources allocated at time t, denoted as *R(t)*. Specifically, *σ(t)* is modeled as a function of resources *R(t)*:

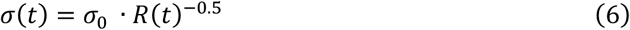

Here, *σ_0_* is a scaling factor defined as the standard deviation of the simulated variable (e.g., position or angle) across all stimuli. This formulation ensures that greater resource allocation leads to lower simulation noise. Resource allocation *R(t)* is dynamically optimized by minimizing an objective function that balances prediction accuracy against resource expenditure.

### Definitions

*S_t_*: Scene state vector at time *t*, formally defined as:

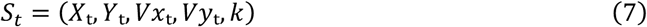

Where (*Xₜ*, *Yₜ*) are Cartesian coordinates of the blue ball’s centroid, (*Vxₜ*, *Vyₜ*) are the blue ball’s instantaneous velocity components, *k* is the mass ratio between the two balls. We didn’t include the kinematic variables of the white ball in the state vector, because: 1) the white ball remains stationary at a fixed location pre-collision; 2) its post-collision trajectory does not affect the computation of the blue ball’s exit point required for the experimental task.

φ: State transition function governed by Newtonian mechanics, such that:

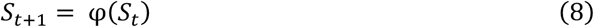

### The Objective Function

As defined in the main text, the total cost *C_total_* (Eq. 2) is composed of a behavioral cost term (Eq. 3) and a resource cost term (Eq. 4):

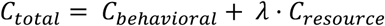

Where:

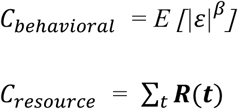

Here, the behavioral cost *C_behavioral_* captures the prediction error *ε* through a power-law relationship^43^, and the resource cost *C_resource_* represents the total computational resources allocated over time. The weighting factor *λ* determines the relative importance between behavioral and resource costs, allowing the model to flexibly prioritize either prediction accuracy or computational efficiency based on the desired trade-off. The parameter *β* controls the transformation of estimation error *ε* into the behavioral cost through a single-parameter power-law relationship. Resource allocation is dynamically optimized for fixed *λ* and *β* by minimizing this objective function using Particle Swarm Optimization (PSO)^73^, implemented via the particleswarm function in the MATLAB Global Optimization Toolbox (R2024a, The MathWorks, Inc., Natick, MA, USA)^74^. The PSO parameters were set as follows: SwarmSize = 400, MaxIterations = 500, FunctionTolerance = 1×10⁻³.

### Model fitting

To fit the model’s free parameters (*λ* and *β*) to human behavioral data, we implemented a nested optimization procedure composed of two interleaved loops. The outer loop iteratively derives the parameter pair (λ, β) that maximizes behavioral fit. For each candidate (λ, β) evaluated by the outer loop, an inner optimization dynamically allocates cognitive resources across the simulation timeline. This inner loop minimizes the total cost by balancing prediction accuracy against resource expenditure, optimizing the resource trajectory for given (λ, β) values. Using this optimized resource allocation, the model generates predicted distributions of the blue ball’s exit angle, which the outer loop compares with behavioral data to update *λ* and *β*. This bi-level architecture yields human behavior-consistent predictions while maintaining a principled account of cognitive cost.

Specifically, we fitted the model to behavioral data in the outer loop using Bayesian optimization^44^ to find the parameter vector *θ* = {*λ*, *β*} that maximizes the log likelihood function:

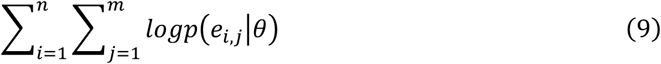

where *e_i_*_,*j*_ denotes participant *j*’s estimation error of the blue ball’s exit angle in trial *i*, and *p*(*e_i_*_,*j*_|*θ*) represents the probability density of observing this error under model parameter *θ*. Bayesian optimization is well suited here because the log likelihood in our model is evaluated via Monte Carlo simulations with stochastic noise, resulting in a non-smooth objective for which gradients are unavailable.

The model’s predicted estimation error distribution for a given parameter vector *θ* was generated as:

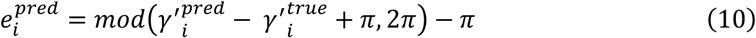

where 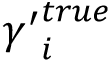 is the ground-truth exit angle in trial *i*, derived through Newtonian mechanics, and 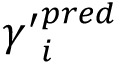 is a vector of predicted exit angles generated by Monte Carlo simulation for trial *i*, with the sample size corresponding to the Monte Carlo simulation count (*n* = 100). The modulo operation ensures that the angular errors remain bounded within [−π, π], accounting for the circular nature of angle estimation.

To obtain a continuous probability density function over errors, we applied circular kernel density estimation (KDE) to these samples. The log-likelihood for a given participant’s response was then computed by evaluating the KDE at the observed error *e_i_*_,*j*_.

Crucially, the resource allocations *R(t)* are not free parameters for fitting behavioral data, but are dynamically optimized by the model to minimize *C_total_* under fixed *λ* and *β*. Therefore, the total number of free parameters in this model remains two.

### Fixed resource IPE model Overview

The fixed resource IPE shares the same foundation as the dynamic resource IPE, differing only in its treatment of internal simulation noise: here, the noise level is held constant across time and experimental conditions. Under the assumption of a power-law relationship between noise and resources, the resource value (*R*_0_) is likewise constant over time and is fitted as a free parameter during model estimation.

### Model fitting

The model contains one free parameter, *R*_0_. We fitted the model to behavioral data using Bayesian optimization^44^ to find the parameter *R*_0_ that maximizes the log-likelihood function:

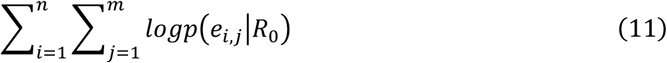

where *e_i_*_,*j*_denotes participant j’s estimation error of the blue ball’s exit angle in trial *i*.

The predicted estimation error distribution for this model was computed using the same approach as described for the dynamic resource IPE model. Specifically, for each trial, the fixed resource IPE model generated a set of predicted exit angles through Monte Carlo simulation under the parameter *R_0_*. The estimation errors were then obtained by subtracting the ground-truth exit angle 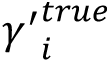 from each predicted sample 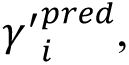 followed by a modulo operation to confine errors within the [−π, π]. Probability densities were estimated through circular KDE, with log-likelihoods derived by evaluating the KDE at each participant’s observed error *e_i_*_,*j*_.

### Extended perceptual-simulation dynamic resource IPE model

To examine whether the resource allocation mechanism generalizes to conditions in which perceptual input is available during the visible period, we extended the main IPE framework by incorporating both perceptual updating and internal simulation. As in the original IPE framework, the extended model was implemented in both dynamic-resource and fixed-resource forms. Here we first describe the extended dynamic-resource version.

In the original dynamic resource IPE model, uncertainty arises entirely from internal simulation noise, and the entire trajectory is treated as a forward simulation process beginning from the initial state. In contrast, the extended model distinguishes between visible and invisible epochs. During the visible epoch, state estimates are updated from the true scene state with added perceptual noise, whereas during the invisible epoch, they are propagated through internal forward simulation with simulation noise.

Specifically, during the visible epoch, the state estimate at time *t* is given by:

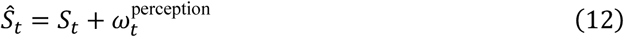

where the perceptual noise is modeled as a zero-mean Gaussian distribution:

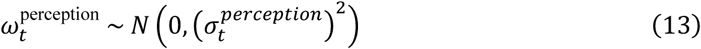

After occlusion onset, the model switches to internal simulation dynamics:

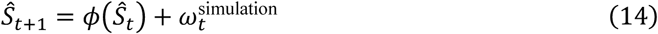

where the simulation noise is likewise modeled as a zero-mean Gaussian:

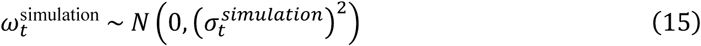

In both stages, noise magnitude is modulated by dynamically allocated resources *R(t)*. The simulation noise term follows the same resource-dependent formulation as in the main dynamic resource IPE model:

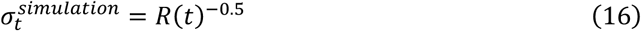

To characterize the relative reliability of perceptual updating and internal simulation, we introduced a perception-simulation noise ratio parameter, *k_noise_*, such that the perceptual noise term was defined as:

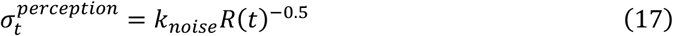

Thus, *k_noise_* controls the relative magnitude of perceptual and simulation noise under the same resource level. We evaluated the model under three values of *k_noise_* (0.25, 0.5, and 1) to systematically probe different regimes of perceptual–simulation noise balance.

All other aspects of the model, including the objective function, the dynamic resource optimization procedure, Monte Carlo sampling, and behavioral fitting framework, were identical to those of the main dynamic resource IPE model.

### Extended perceptual-simulation fixed resource IPE model

Analogous to the dynamic-resource version, we also implemented an extended perceptual-simulation fixed resource IPE model. During the visible epoch, state estimates were updated from the true scene state with added perceptual noise, whereas during the invisible epoch, they evolved through internal forward simulation with simulation noise.

Perceptual and simulation noise followed the same power-law relationship with resources as in the extended dynamic-resource model, with their relative magnitudes controlled by the perception-simulation noise ratio parameter *k_noise_*:

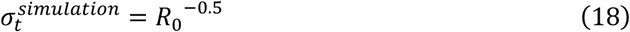

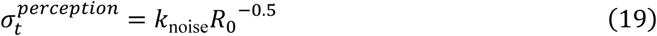

Unlike in the dynamic-resource version, the resource level remained fixed across time and conditions and was fitted as the sole free parameter. All other aspects of Monte Carlo sampling, likelihood estimation, and behavioral fitting were identical to those described for the main fixed resource IPE model.

## Data availability

All data supporting the current study have been deposited at Zenodo and are publicly available at the time of publication (https://doi.org/10.5281/zenodo.17157343). Source Data are provided with this paper.

## Code availability

The custom codes related to computational modeling have been deposited at Zenodo and are publicly available at the time of publication (https://doi.org/10.5281/zenodo.17157343).

## Acknowledgements

This work was supported by the Strategic Priority Research Program of the Chinese Academy of Sciences (XDB1010101) and Brain Science and Brain-like Intelligence Technology--National Science and Technology Major Project (2022ZD0204600); We thank the 3T core facility of CAS CEBSIT/ION for technical support; We thank the MEG facility of CAS CEBSIT/ION for the assistance in data collection; We thank the members of the Chang lab for helpful comments on the manuscript.

## Author contributions

L.L. and Q.W. performed the experiments; L.L. analyzed the data and constructed the computational models; L.L. and L.C. conceived the project; Q.Y. and L.C. supervised the project; L.L. and L.C. wrote the initial draft; all authors revised the manuscript and approved the final version.

