## Supplementary figures for "Dynamic resource allocation orchestrates physical simulation in the human brain"

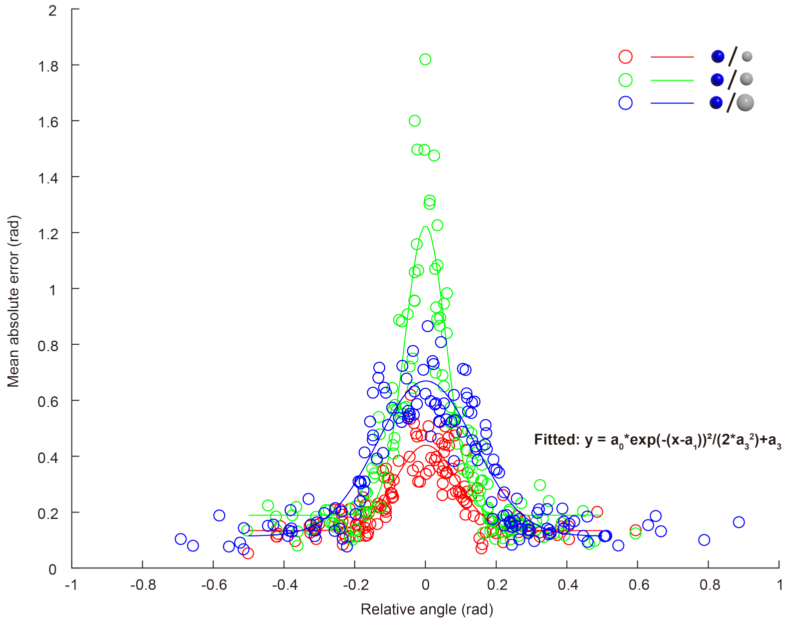


**Figure S1. Relationship between participants' prediction errors and relational physical variables**

Mean absolute error across all participants per trial is plotted against the relative angle $\alpha$ (open circles). Curves show Gaussian-function fits with scaling and offset parameters. Colors represent mass ratio $k$: red indicates trials where the blue ball is heavier than the white ball; green indicates equal mass; and blue indicates trials where the blue ball is lighter than the white ball.

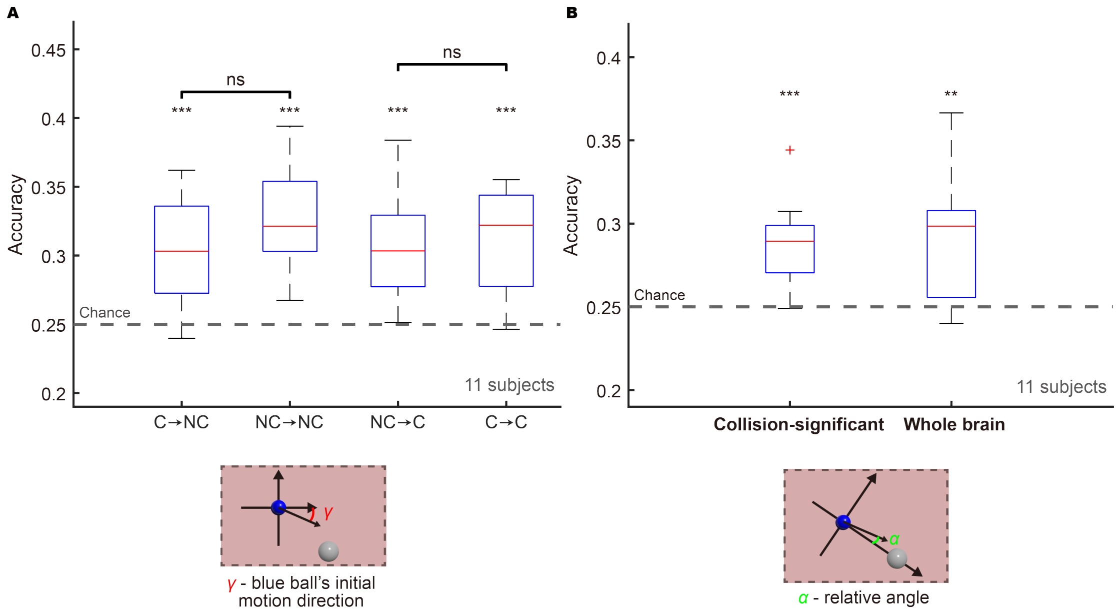


**Figure S2. Multivariate decoding of physical variables using fMRI data**

**(A)** Multivariate decoding of the blue ball's initial motion direction ($\gamma$) using cross-trial-type and within-trial-type schemes. For cross-trial-type decoding (e.g., C⟶NC), classifiers were trained on collision trials (C) and tested on non-collision trials (NC), or trained on non-collision trials and tested on collision trials (NC⟶C). For within-trial-type decoding (C⟶C and NC⟶NC), classification was performed separately within each condition using five-fold cross-validation. Box plots show the median and interquartile range. Chance level was 25% (four-class classification). Asterisks indicate significant above-chance decoding (*** = $p<0.001$, one-tailed Student's t-test); n.s. indicates no significant difference between decoding schemes (two-tailed Student's t-test).

**(B)** Multivariate decoding of relative angle ($\alpha$) in non-collision trials. The left panel shows decoding using vertices significantly tuned to relative angle in collision trials (identified via univariate ANOVA), whereas the right panel shows decoding using all vertices across the whole brain. Decoding accuracy was estimated using five-fold stratified cross-validation. Box plots show the median and interquartile range. Chance level was 25% (four-class classification). Asterisks indicate significant above-chance decoding (** = $p<0.01$; *** = $p<0.001$, one-tailed Student's t-test).

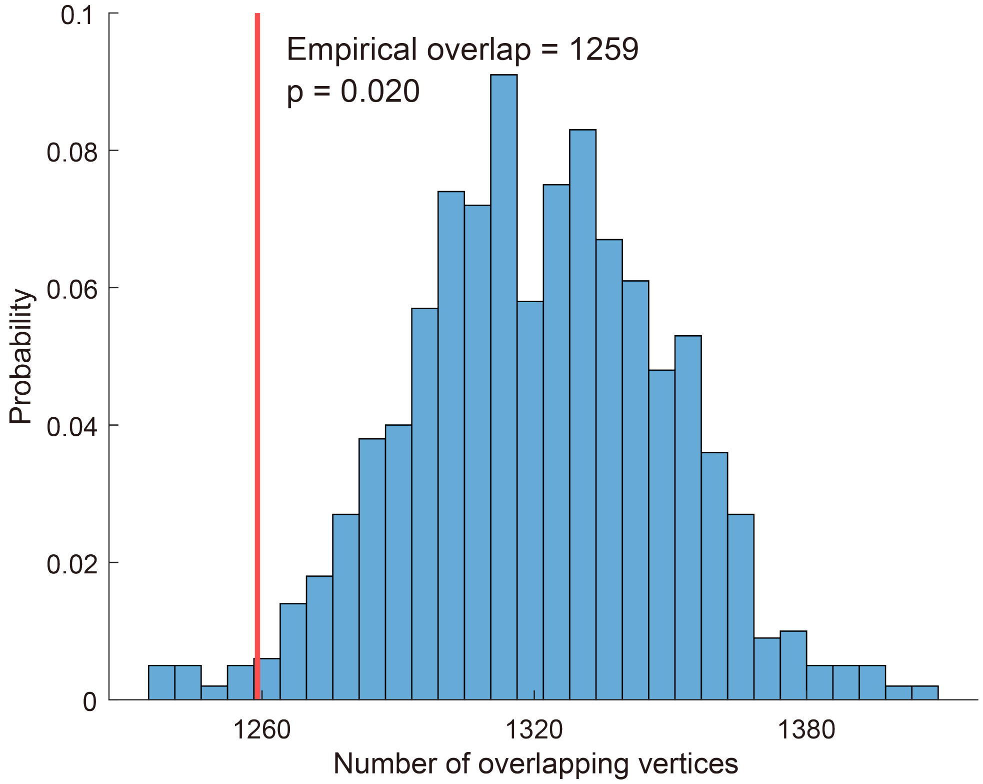


**Figure S3. Permutation-based test of spatial overlap across physical variables**

Null distribution of multi-variable spatial overlap among the four physical variables shown in Fig. 3D ($\gamma$, $\gamma'$, $\alpha$, and $k$). Spatial overlap was quantified as the number of vertices selective for two or more physical variables. The null distribution was generated by randomly shuffling the cortical locations of variable-selective vertices for each variable across the cortex while preserving the number of selective vertices per variable (1,000 permutations). The observed overlap (vertical line) was significantly lower than expected by chance ($p = 0.020$).

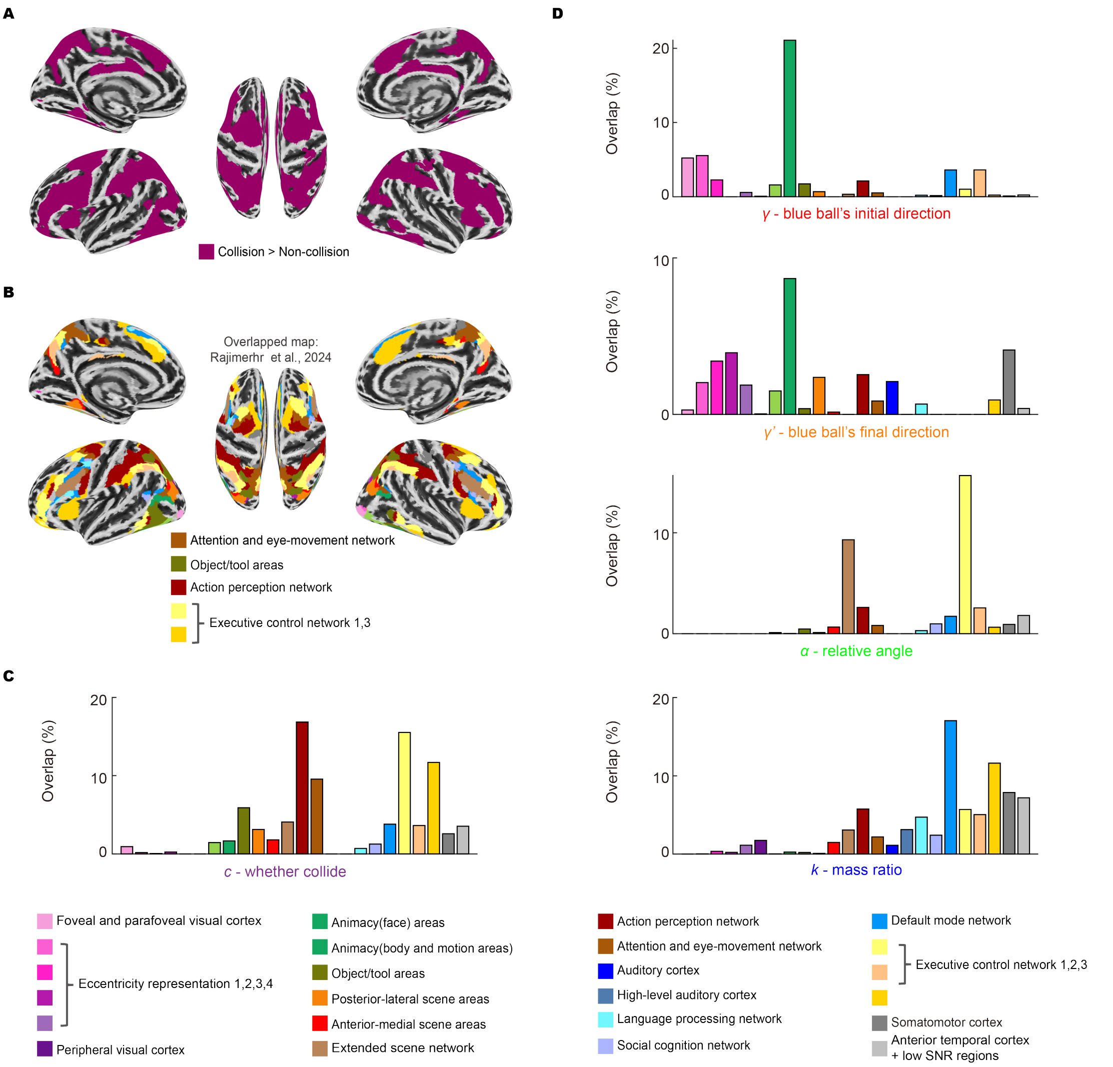


**Figure S4. Overlap between brain regions representing physical variables and the naturalistic movie-derived parcellation.**

**(A)** The purple regions indicate brain areas showing significantly stronger activation during collision trials than during non-collision trials (11 participants, FDR corrected, $p < 0.05$).

**(B)** The collision-activated region identified in (A) overlaid on the naturalistic movie-derived parcellation (only clusters with a Jaccard index > 0.05 are shown).

**(C)** Spatial overlap between collision-activated regions and the 24 movie-derived functional networks, without applying a Jaccard index threshold (cf. Fig. 3F).

**(D)** Spatial overlap between brain regions exhibiting significant tuning to physical variables ($\gamma$, $\gamma'$, $\alpha$, and $k$) and the 24 movie-derived functional networks, without applying a Jaccard index threshold.

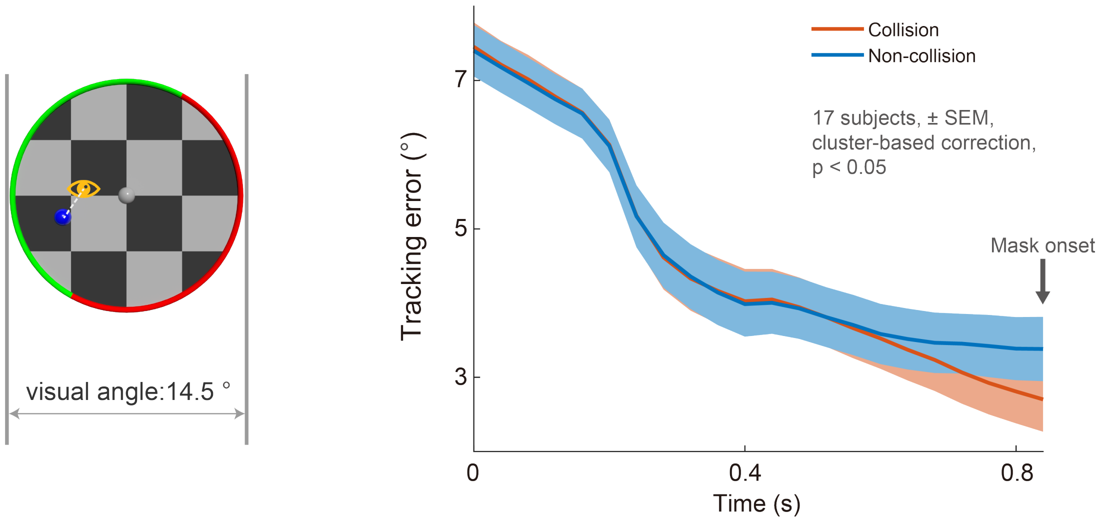


**Figure S5. Gaze tracking error dynamics in collision and non-collision trials**

Gaze tracking error was defined as the Euclidean distance, in degrees of visual angle, between the gaze position and the instantaneous center of the blue ball during the visible epoch. Tracking error was computed at each time point and then averaged across trials separately for collision and non-collision trials. Curves show the group mean across participants, and shaded regions indicate SEM. Vertical arrow indicates mask onset.

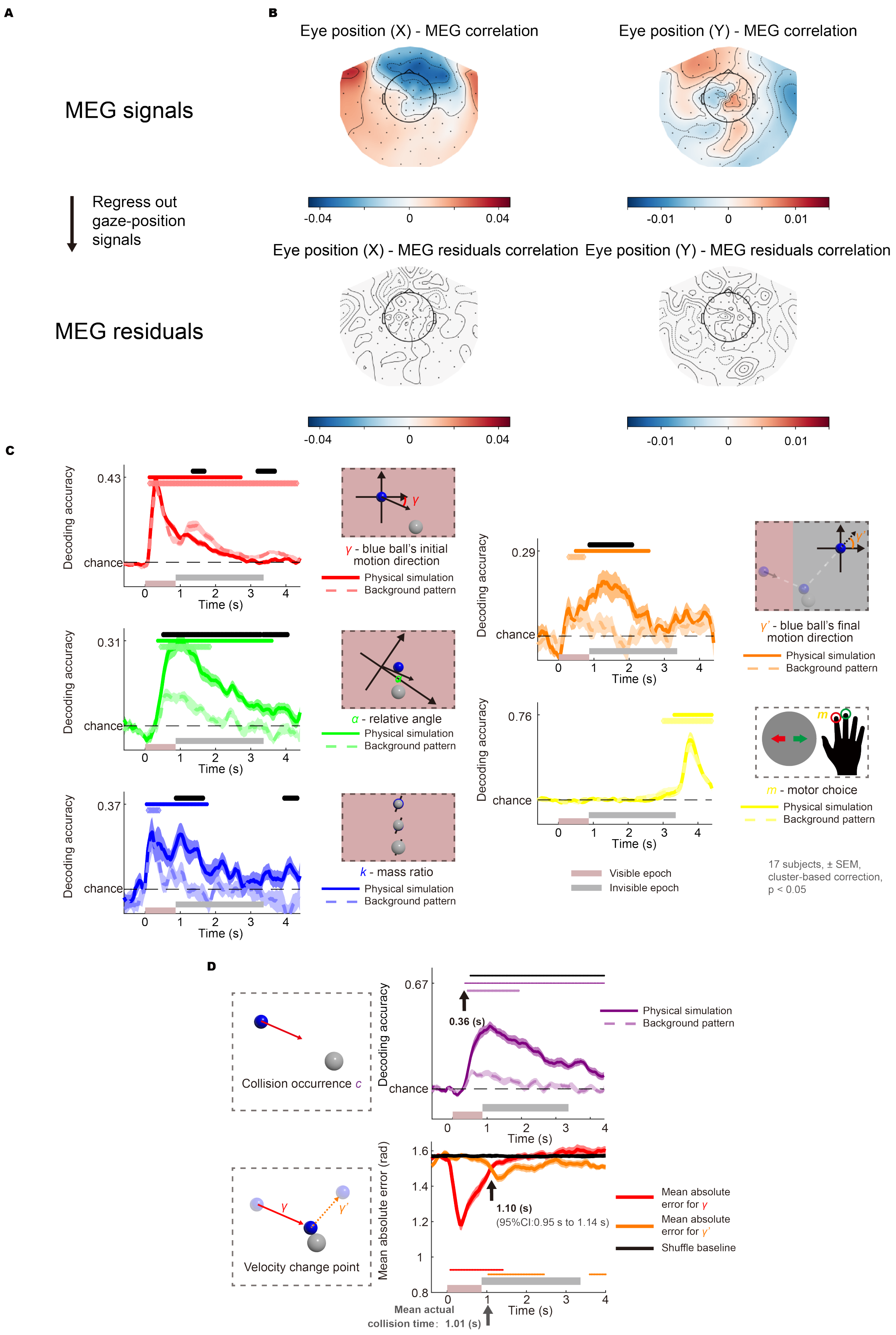


**Figure S6. MEG results after regressing out eye-movement signals.**

Replication of analyses in Fig. 4B-D and Fig. 5A using eye-movement-corrected MEG signals. All procedures were identical to those in the main text, except that horizontal and vertical eye positions were first regressed out from the raw MEG signals at the single-subject level, and the resulting residuals were used for all subsequent analyses.

(A) Schematic illustration of the eye-movement regression procedure. At each time point and for each participant, trial-wise horizontal and vertical gaze positions were included as linear regressors and regressed out from the MEG signals.

(B) Correlation between MEG signals and eye position before and after regression. Raw MEG signals showed systematic correlations with eye position, which were effectively removed in the residual signals, confirming successful gaze-position regression.

(C-D) Replication of the main analyses in Fig. 4B-D and Fig. 5A using MEG residuals. All key temporal patterns were preserved after controlling for eye movements.

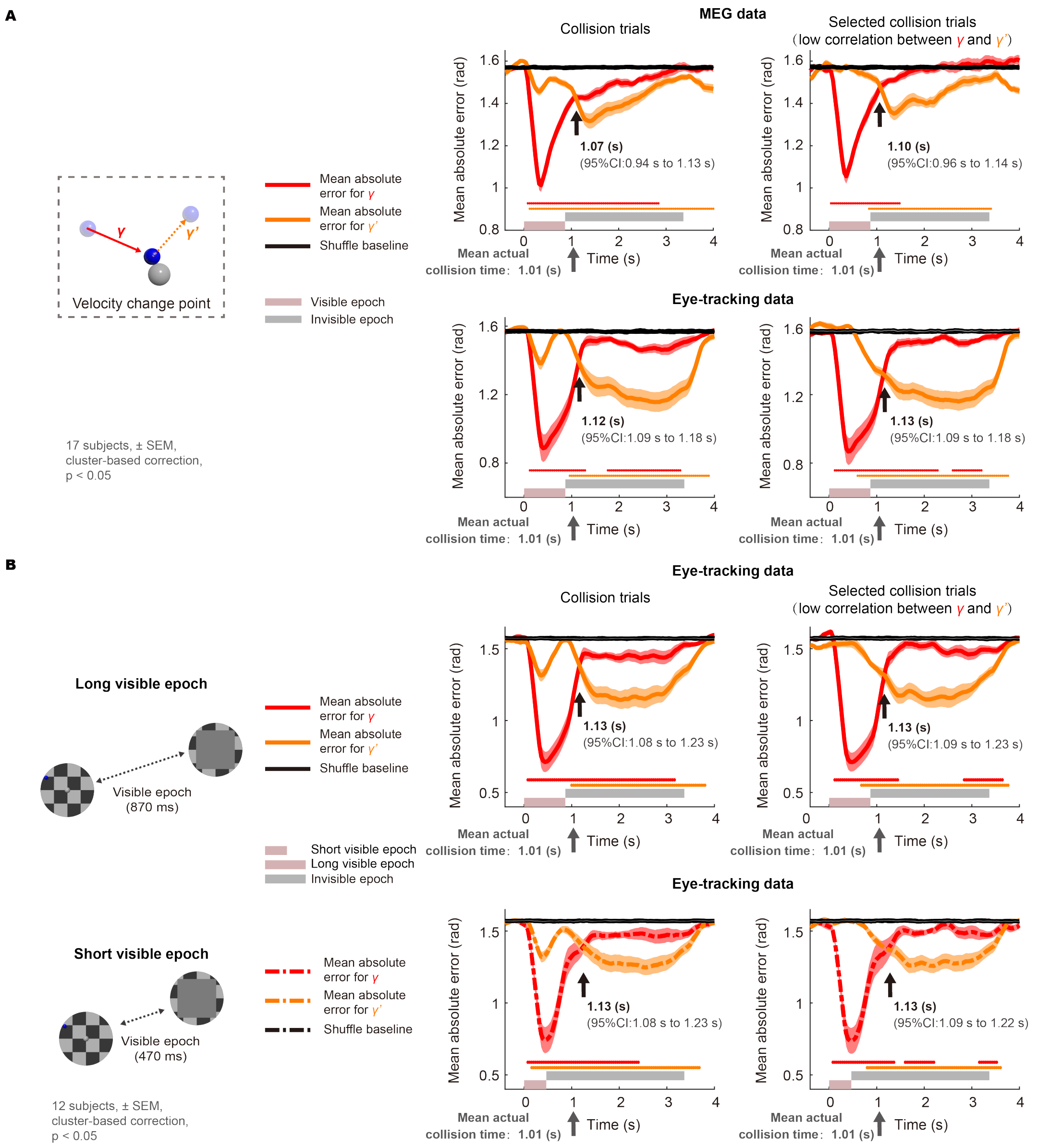


**Figure S7. Real-time simulation of blue ball motion direction is reflected in both neural activity and eye-tracking data**

**(A)** Same convention as Fig. 5A (bottom panel). Decoding analyses employ ridge regression, with decoding errors quantified as the mean absolute angular difference between predicted and actual directions ($n$ = 17 participants, $p$ < 0.05, light-shaded region indicates SEM, cluster-based correction). The top and bottom rows show decoding results based on MEG data and eye-tracking data, respectively. The middle and right columns display all collision trials and a subset of collision trials with minimized correlation between $\gamma$ and $\gamma'$, respectively (see Methods).

**(B)** Same convention as Fig. 5C (third column). Decoding analyses employ ridge regression, with decoding errors quantified as the mean absolute angular difference between predicted and actual directions ($n$ = 17 participants, $p$ < 0.05, light-shaded region indicates SEM, cluster-based correction). Both rows show decoding results based on eye-tracking data. The middle and right columns display all collision trials and a subset of collision trials with minimized correlation between $\gamma$ and $\gamma'$, respectively (see Methods).

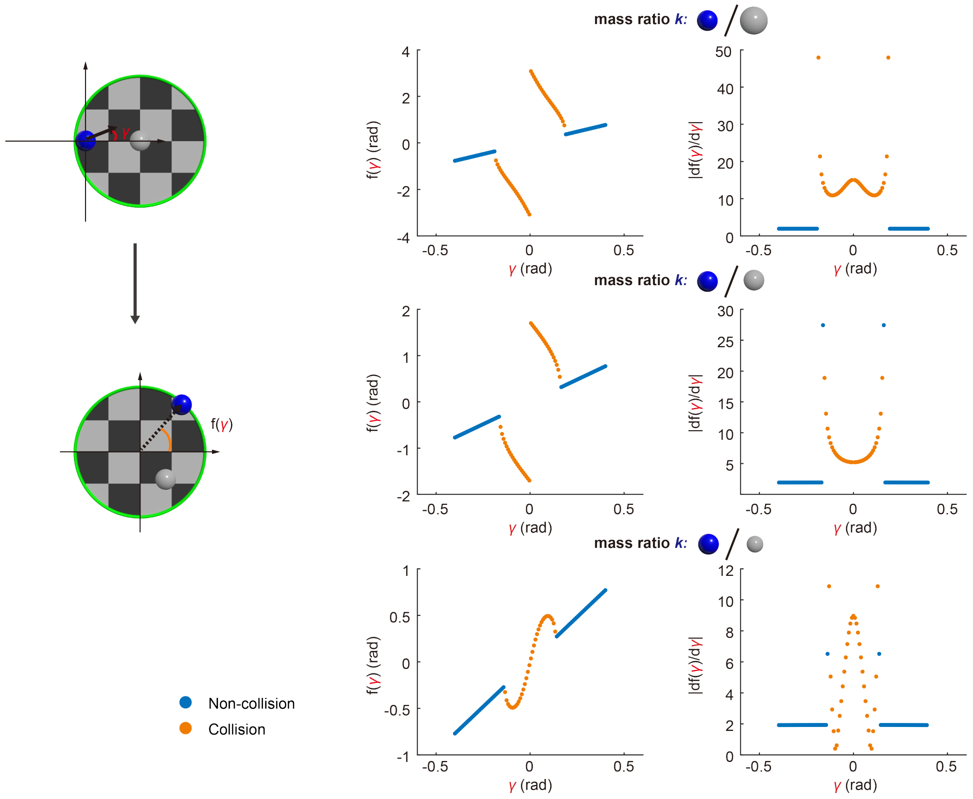


**Figure S8. Impact of perturbations in the blue ball's initial motion direction on its final exit point.**

Left column: $\gamma$ denotes the blue ball's initial motion direction, and f($\gamma$) denotes its exit angle from the disc. For visualization purposes, the blue ball's initial position is fixed at the leftmost point of the disc.

Middle column: Relationship between the blue ball's initial motion direction $\gamma$ and its exit angle f($\gamma$), with each row corresponding to a different mass ratio.

Right column: Relationship between df($\gamma$)/d$\gamma$ and $\gamma$ under different mass ratios. Higher derivatives in collision trials indicate greater sensitivity to initial directions ($\gamma$) perturbation.

In all panels, orange indicates collision trials, and blue denotes non-collision trials.

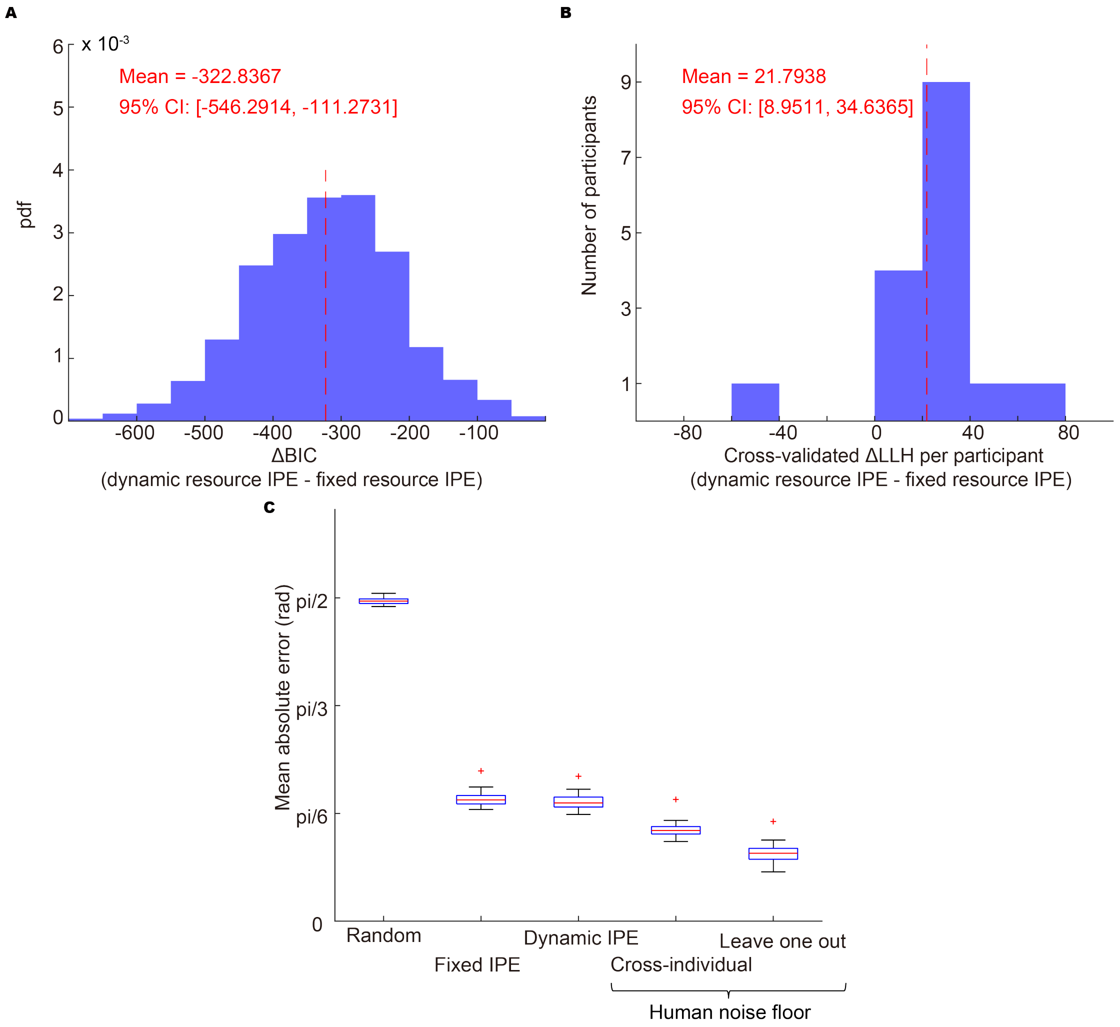


**Figure S9. Additional analyses of the dynamic and fixed resource models.**

**(A)** Distribution of ΔBIC values (dynamic resource IPE – fixed resource IPE), obtained via a bootstrap procedure (1,000 iterations). Negative ΔBIC values indicate better performance of the dynamic resource IPE. The red dashed line indicates the mean ΔBIC across bootstrap iterations.

**(B)** Distribution of ΔLLH values per participant (dynamic resource IPE – fixed resource IPE), based on 4-fold cross-validation, in which 16 participants were divided into four folds of four participants each. In each iteration, one fold served as the test set and the remaining three folds (12 participants) served as the training set. Each participant thus contributed one ΔLLH value when serving in the test set. Positive ΔLLH values indicate better performance of the dynamic resource IPE. The red dashed line indicates the mean ΔLLH across participants.

**(C)** Comparison of MAE between model predictions and behavioral responses across four conditions: random baseline, fixed resource IPE, dynamic resource IPE, and human noise floor. MAE was computed for each subject. For the two IPE models, each trial yielded a distribution of stochastic predictions (100 samples per trial); errors were computed for each sample and averaged across simulations and trials. The random baseline was generated by uniform sampling from [−π, π]. Human-level predictability was estimated using two measures: cross-individual human-human consistency and leave-one-out (LOO) consistency. The cross-individual measure treats other participants' responses as samples from the human response distribution and computes circular absolute error between each sample and the corresponding participant response, similar to our treatment for IPE models, whereas the LOO measure compares each participant with the circular mean of the remaining participants and therefore provides a lower-variance, more stringent estimate of the human noise floor. Wilcoxon signed-rank tests showed that dynamic resource IPE had lower MAE than fixed resource IPE ($p < 0.001$), both IPE models outperformed the random baseline ($p < 0.001$), and both human-level baselines (cross-individual and LOO) exhibited lower MAE than either IPE model ($p < 0.001$).

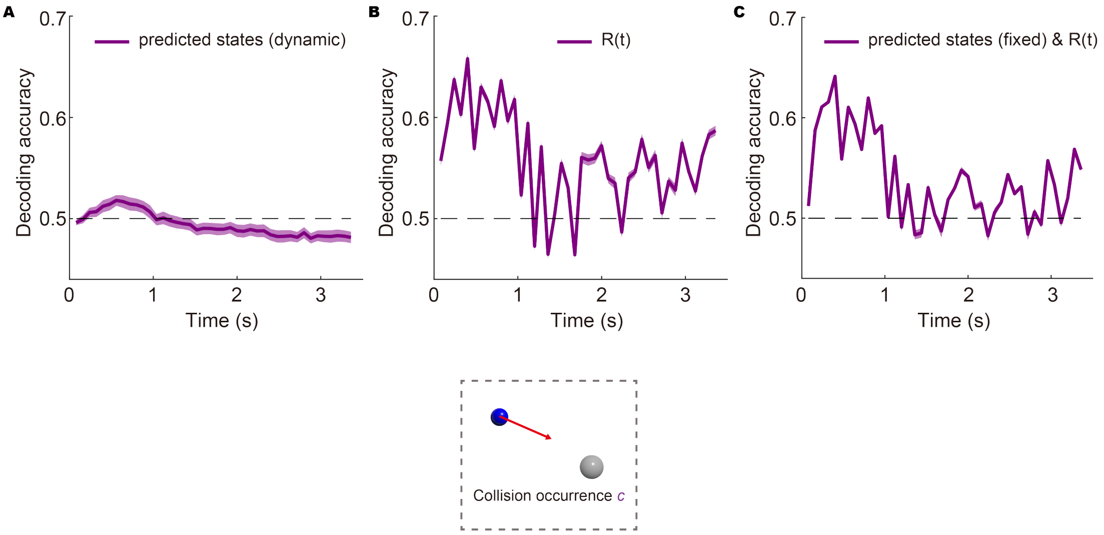


**Figure S10. Model-component ablation analyses of collision-occurrence decoding.**
Decoding accuracy for collision occurrence was computed using a binary SVM. Ablation analyses were performed to disentangle the contributions of different model components:
(A) Decoding based on predicted states alone in the dynamic resource model, excluding *R(t)*;
(B) Decoding based on *R(t)* alone;
(C) Decoding based on predicted states in the fixed resource model augmented with *R(t)* from the dynamic resource model.

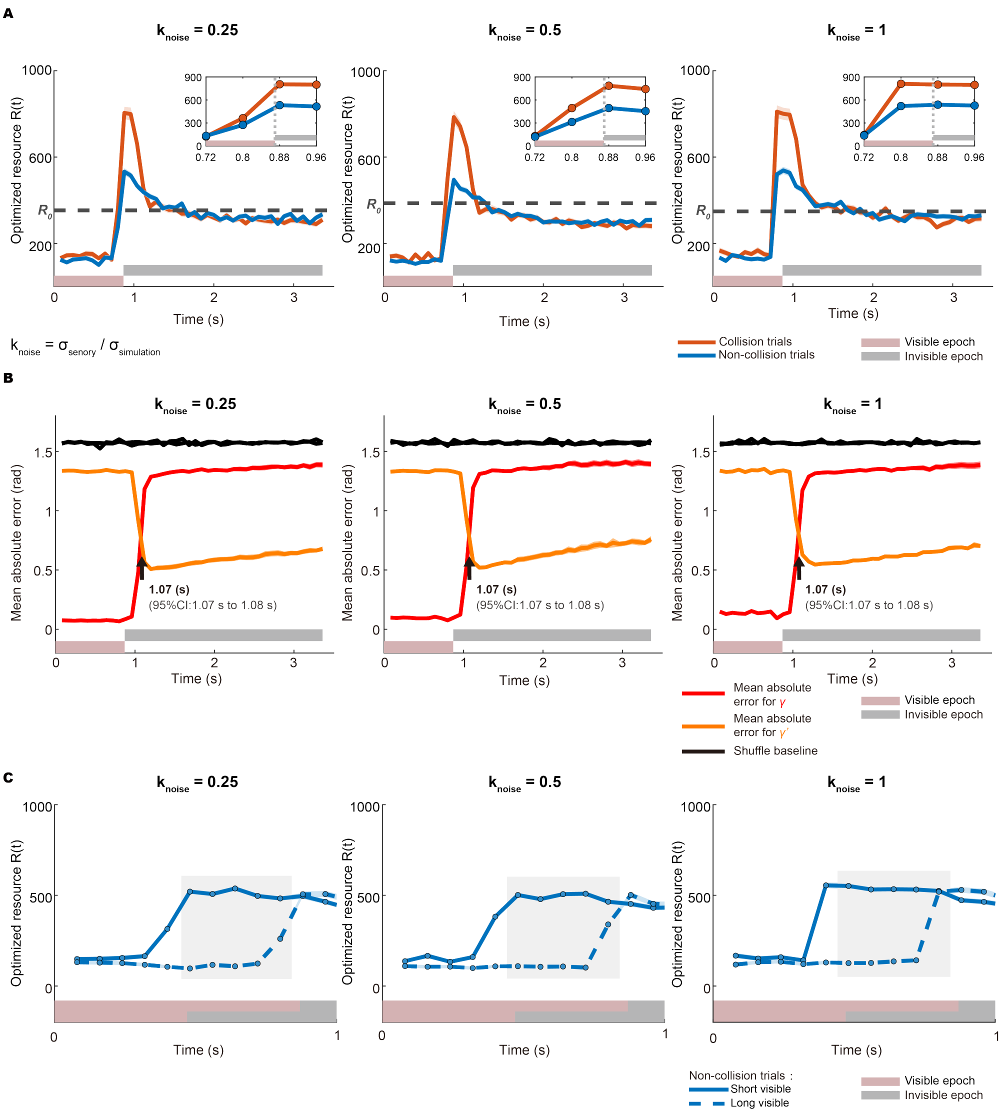


**Figure S11. Extended perceptual-simulation IPE model**

(A) Time course of single-trial resource allocation estimates obtained by optimizing the extended perceptual-simulation IPE model using the best-fitting parameters, averaged separately for collision and non-collision trials. Shaded areas indicate SEM. Dashed horizontal lines denote the fitted constant resource level $R_0$ from the fixed resource IPE model, serving as a reference. Each column shows results for a different value of the perception-to-simulation noise-scaling parameter $k_{noise}$. Insets show magnified views around occlusion onset at 0.87 s; dashed vertical lines indicate occlusion onset.

(B) Decoding analysis used to estimate the model-implied collision time. The horizontal and vertical components of the blue ball's pre- and post-collision motion directions were decoded using ridge regression. The decoding features consisted of the predicted states and the resource allocation term *R(t)* derived from the extended perceptual-simulation IPE model under different $k_{noise}$ settings. Decoding errors were quantified as mean absolute angular differences between predicted and actual directions. The decoding error curves for the motion direction of the blue ball before ($\gamma$) and after ($\gamma'$) the collision consistently intersected at approximately 1.07 s across $k_{noise}$ values, with the 95% confidence interval estimated by bootstrapping.

(C) Time course of single-trial resource allocation estimates obtained by optimizing the extended perceptual-simulation IPE model using the same hyperparameters, with two different stimulus visibility durations (long = 0.87 s vs. short = 0.47s). Only non-collision trials were included. Shaded rectangles highlight identical time points in the physical trajectory across trials, which correspond to visible segments in the long-duration condition but occluded segments in the short-duration condition, allowing direct comparison of resource allocation under different visibility.

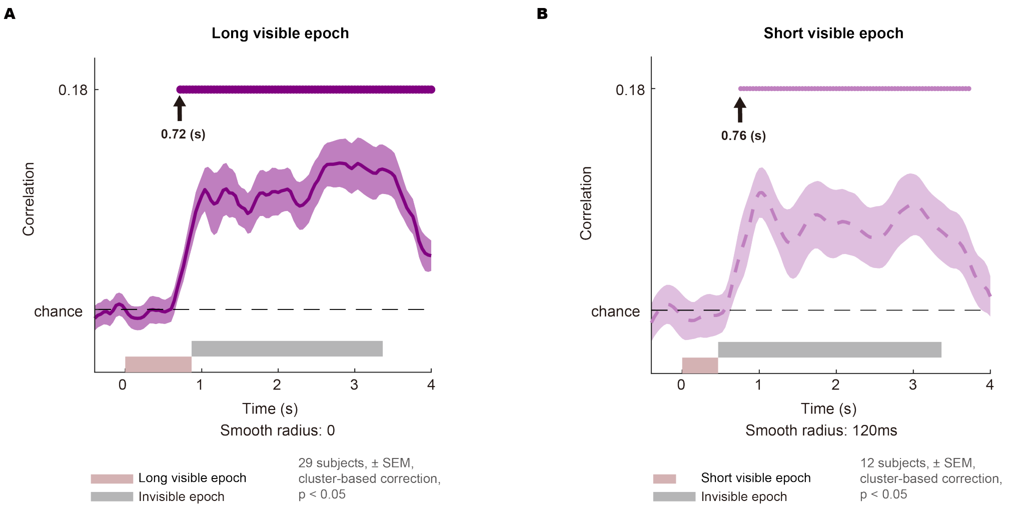


**Figure S12. Additional pupil size analyses.**

**(A)** Unsmoothed correlation analysis between pupil size and collision occurrence. While Fig. 6E applied a 120 ms smoothing window to the correlation results (which potentially advanced the apparent timing of significant positive correlations), this unsmoothed analysis confirms the original estimate of 0.72s.

**(B)** Same as Fig. 6E, but for the short visible epoch condition. Positive correlations began at a comparably early time (0.76 s).

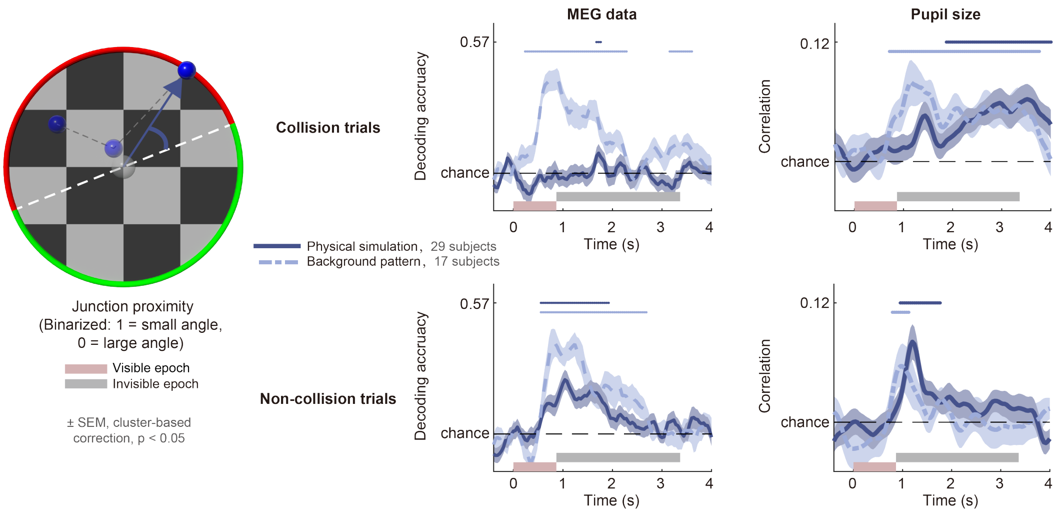


**Figure S13. Representation of junction proximity in collision and non-collision trials.**
Same analysis as in Fig. 6F, but conducted separately for collision and non-collision trials, with the background pattern condition also included. Notably, in physical simulation conditions, junction proximity signal is stronger and faster in non-collision trials compared to collision trials. This likely reflects resource allocation limits: when resources are assigned to collision simulations, it becomes difficult to allocate additional resources to the subset involving junction proximity within collision trials. Future research should clarify how resources are distributed between the two types of sensitivity-critical variables.

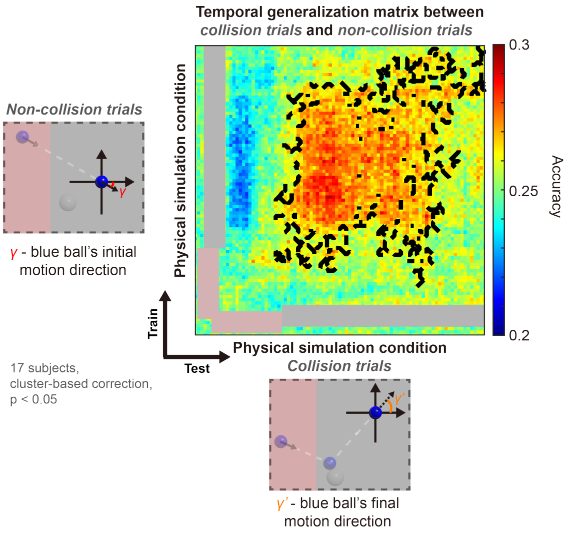


**Figure S14. Generalizable decoding of simulated motion direction between collision and non-collision trials.**

365 Same convention as Fig. 7C, but here the $\gamma$-direction decoder is trained using non-collision trials of the physical simulation condition and tested to predict $\gamma'$ in collision trials.
